# Comparing re-referencing methods for human intracranial EEG

**DOI:** 10.64898/2026.10.13.758866

**Authors:** Luca D. Kolibius, George Parish, Marije ter Wal, Mircea van der Plas, David T. Rollings, Stephanie Gollwitzer, Katrin Walther, Rüdiger Hopfengärtner, Hajo Hamer, Maria Wimber, Simon Hanslmayr

## Abstract

Stereo-electroencephalography (SEEG) involves the implantation of depth electrodes in the brain to monitor epileptic seizures for surgical feasibility. As the electrodes advance into deeper limbic structures, the macro contacts distributed along the shaft record electrical activity from more lateral sites, while a bundled array of microwires simultaneously captures highly localised signals from the tip of the electrode. In a clinical setting these electrical signals are recorded relative to a reference signal, which may be sub-optimal for subsequent analyses that are unrelated to clinical objectives. Therefore, a secondary re-referencing operation is often undertaken to transform the signal in relation to a specific hypothesis. Despite a recent surge in SEEG studies, comparative studies on the much-understudied issue of re-referencing are sparse. Here, we analyse data collected from 16 human patients during an episodic memory task and comparatively apply a range of re-referencing methods to both macro contact and micro-wire data. We find that the choice of method changes the proportion of trials that survive artefact rejection, low frequency power, and the aperiodic background against which that power is read. In the hippocampal recordings, however, no method differed from any other in the size of a memory effect in power or spike–field coherence. This early transformative decision therefore bears on how the signal is described. We found no evidence that it changes the detectability of the behavioural effects tested here, and no method reversed the direction of a memory effect.

## 1. Introduction

Stereo-electroencephalography (SEEG) involves the implantation of depth electrodes for the purpose of monitoring epileptic seizures. As SEEG offers the prospect of recording electrical signals in the brain at both a high spatial and temporal resolution, there has been a recent drive to recruit epileptic patients for human brain research. The modes of data acquisition that SEEG offers also differ in specificity. Macro contacts down the length of an implanted electrode offer the ability to record the fluctuations of electrical fields that are generated by the summated activity of large groups of neurons (Schomer & Lopes da Silva, 2018). Microwires can also be inserted inside the hollow tube of a depth electrode, splaying out from the tip to record highly localised activity down to the single neuron level (Suthana & Fried 2012).

Electrical signals can best be described as a dynamic measurement of potential differences between two points (Nunez & Srinivasan, 2006). Therefore, at the point of data acquisition target electrical signals must be measured in reference to another signal of choice. However, these reference signals are rarely inactive (Mercier et al., 2017), which can have two effects. First, the target signal could be contaminated by a different neural signal which is volume conducted to the reference electrode (Herreras, 2016). For instance, alpha oscillations in the neocortex may be picked up by reference electrode on the mastoid and consequently contaminate signals recorded from a hippocampal contact. The same applies to non-neural signals such as muscle activity or power line noise which may be picked up by the reference contact. Therefore, an additional re-referencing step is often undertaken in an attempt to clean up the signal and/or improve the local resolution.

There are many re-referencing strategies available that each transform the target signal in very different ways (see Figure 1). For example, common noise might be reduced by removing a relatively inactive source (such as one in a white matter or subdural location), or a signal shared between neighbours might be reduced by subtracting a neighbouring contact (allowing subsequent analyses to focus on signal that is more local in nature). Though there have been recent attempts to compare outcomes in this transformative pre-processing step (Li, et al., 2018; Liu, et al., 2021), no gold standard exists for how to re-reference an intracranial EEG dataset.

**Figure 1:**
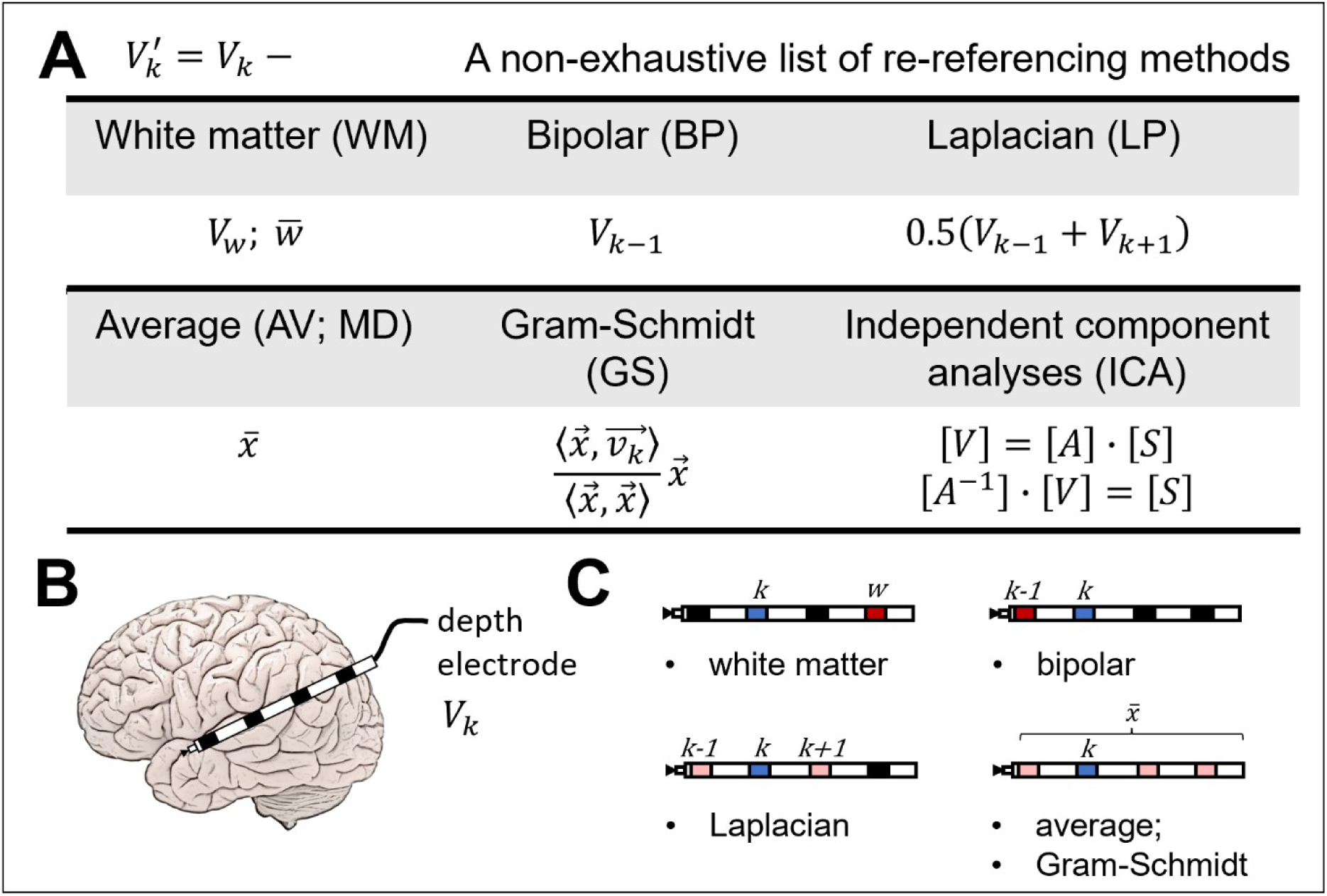
A non-exhaustive list of commonly used re-referencing methods. A: Equations for several re-referencing methods that can be applied to intracranial EEG data. B: Visualisation of a depth electrode. C: Visualisations of the application of the equations in A to the actual data. Blue nodes indicate the contact in question, red nodes indicate single target re-reference contacts and light red indicate a set of re-reference contacts that are averaged. Note that these visualisations do not depict possible re-reference choices that are exterior to the brain, for example in the case of the monopolar mastoid reference, nor do they depict the possibility to average over all depth electrodes within the brain, for example in the case of the average or Gram-Schmidt methods.

SEEG also allows local spiking activity to be analysed together with the field potentials sampled along the same electrode, which has proved a boon for the study of human brain function. The target location of depth electrodes is often the medial temporal lobe (MTL), which provides necessary functions relevant to memory (Eichenbaum, et al., 2007; Squire, et al., 2004). Specifically, low frequency theta oscillations (2-8 Hz) have long been thought to play a formative role in these memory operations (Buzsaki, 2002), and both increases and decreases of theta power with successful memory have been reported. This effect might be made up of a narrowband and a broadband component, and differences in paradigm, recording technique and referencing scheme might emphasise one component over the other (Herweg et al., 2020). The comparisons of re-referencing methods that exist were made during motor tasks, with the isolation of local task-related activity as the criterion, and they favoured the Laplacian reference for broadband gamma and for low frequency oscillations alike (Li et al., 2018; Liu et al., 2021). Low frequency activity in the MTL poses a different problem: the signal of interest is expected to be shared across neighbouring contacts (Buzsáki & Draguhn, 2004), so a method that sharpens spatial resolution may remove part of what is being measured. In the present study, we comparatively evaluate the effect of up to 7 different re-referencing strategies on both macro contact and micro-wire SEEG electrophysiological signals. Data were acquired during an episodic memory paradigm. The time–frequency and spike–field coherence analyses that follow focus on three things: low frequency theta oscillations, their modulation by behaviour, and the aperiodic background against which those oscillations are measured. To foreshadow our findings, the results indicate that there is no one-size-fits-all approach when it comes to re-referencing SEEG data. Instead, the choice of the re-referencing method should be part of a wider strategy that is informed by the specific hypothesis of the research project, as each method will transform the data relative to its own assumptions.

## 2. Methods

### 2.1 List of re-referencing methods

Selected re-referencing methods (see Figure 1) are here categorised in three ways: global methods that extrapolate a hypothetical reference source using the entire set of signals; local methods that identify a single (or subset of) reference source(s) from within the entire set of signals; and data-driven methods that “unmix” data into underlying latent components to remove common sources.

#### 2.1.1. Global methods

Removing the average of the entire set of signals is typically seen as a reasonable approximation of activity introduced by the online reference in scalp EEG (Nunez, 2010; Nunez & Srinivasan, 2006). The underlying reasoning is that the sum of the signals of electrodes equidistantly placed on a sphere will be 0. Obviously, intracranial EEG almost always violates this assumption as electrodes are placed in certain brain areas solely defined by the clinical need of the patient, and thus are not well suited to estimate spherical noise. Second, intracranial EEG contacts measure signal that is more local than the summed potential picked up by scalp EEG contacts (Varela, et al., 2001). A commonly used alternative is therefore to remove an average source on an electrode-by-electrode basis (typically termed electrode shaft resonance or ESR), that is here employed via use of the mean (common average or AV) and median (MD). In addition, we also employ a Gram-Schmidt (GS) operation, that seeks to reduce the likelihood of introducing artefacts to re-referenced signals by performing an orthonormalization operation between a given channel and the average source. This way, only signals that are coupled (i.e. with shared variance) between the reference and target electrode are subtracted (Hipp et al., 2012). Furthermore, as the average is scaled by the inner product of the channel and channel average, this operation can boost smaller amplitude local signal whilst bringing down higher amplitude global signal.

#### 2.1.2. Local methods

Identifying a re-reference source(s) from within the set is very common for intracranial EEG studies, possibly as it represents a cleaner way to remove shared activity that stems from the online reference or global oscillatory signals. The monopolar reference makes use of a source located in a white matter or subdural location, due to the hypothesised likelihood that it contains less signals of neural origin, but a similar degree of non-neural signals (i.e. noise). A challenge here is the accurate identification of white matter locations, which is difficult as they are derived from a not wholly accurate normalisation between the pre- and post-op MRI scans that are used for such localisation. Discrepancies between the real location and ascribed location of an electrode may arise due to the imperfect image processing (i.e. co-registration, normalization, etc.) and also due to displacement of brain tissue by electrodes. Therefore, analysing the signal from these contacts might help to improve this step. Due to neurophysiology, we can assume that a white matter contact should have a lower signal to noise ratio compared to a grey matter contact. Therefore, the current study applies a threshold operation to monopolar re-reference selection (see Figure 2). Candidate white-matter contacts were first identified by visual inspection of the electrode localisation. For each electrode shaft, these candidates were compared with grey-matter contacts from the same shaft using LFP variability and mean wavelet power in the 2-12 Hz and 60-90 Hz ranges. Because theta power is typically more strongly expressed in the medial temporal lobe (Hasselmo, 2012), candidate references were retained only if their 2-12 Hz power was not more than 20% higher than the matched grey-matter contacts.

**Figure 2:**
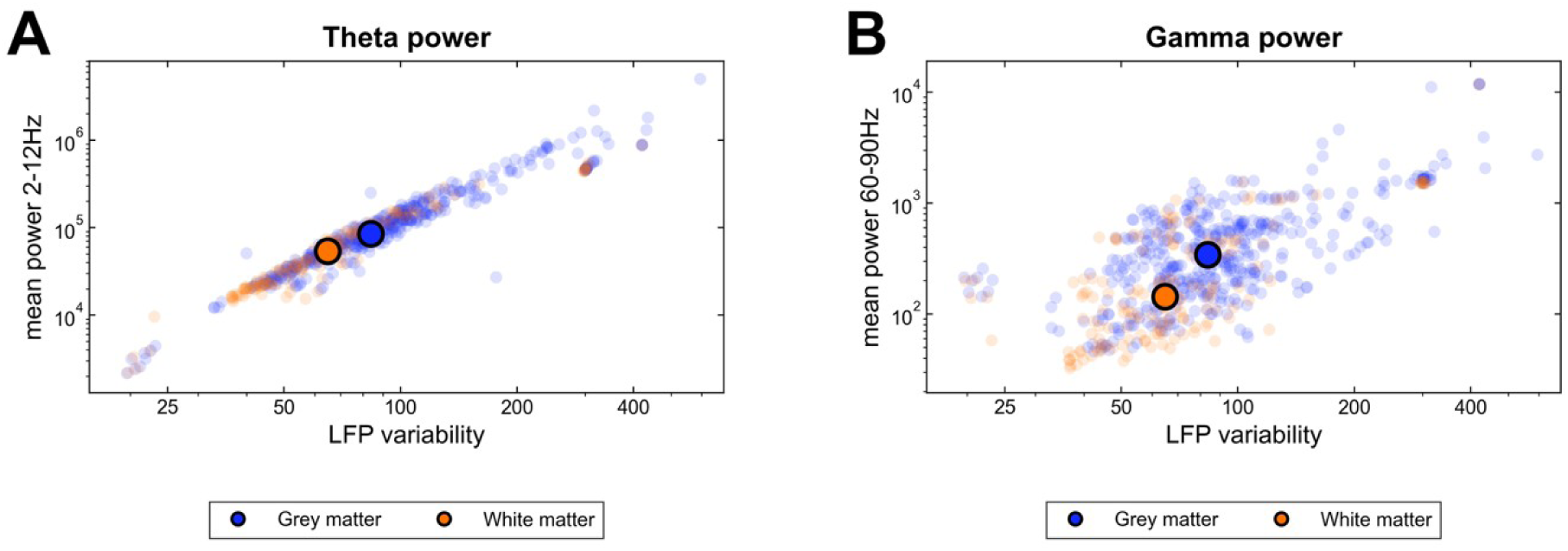
Analyses from an episodic memory paradigm, using SEEG data recorded with depth electrodes, originally referenced using either a bipolar method or a reference contact close to the recording source. A-B: Mean power in the theta (A; 2-12Hz) and gamma (B; 60-90Hz) frequency bands for contacts situated in the grey matter (blue) and white matter (WM; orange), against variability of the local field potential (LFP; calculated as 2 x standard deviation). Blue and orange dots indicate individual grey matter or WM channels, respectively, whilst blue and orange circles mark the median variability and median power of all grey-matter or WM channels, respectively. Both axes use a logarithmic scale.

However, it has been recently shown that the monopolar method might result in the contamination of the signal with task-related signals (Li et al., 2018), causing a high correlation to be introduced across channels. In other comparative studies, neighbour driven methods such as bi-polar and Laplacian were shown to highlight activity that is more local (Li et al., 2018; Liu et al., 2021).

A possible drawback to the use of neighbour driven methods is the loss of information due to the reduced dimensionality of the re-referenced data. For micro-wire data, it is also important to consider that MRI scans are not capable of identifying the exact coordinates of individual wires. As it is thus impossible to identify which wires are neighbours with this kind of data, methods such as bi-polar and Laplacian should instead be considered as randomised mono-polar methods.

#### 2.1.3. Data driven methods

Data driven methods enjoy increasing popularity, due to an increase in computational power and their ability to unmix complex electrophysiological signals which almost always represent a mix of signals from various sources such as local activity, activity from the reference, noise, or volume conducted signals from nearby areas. Independent component analysis (ICA) is one such method, which decomposes the data into its underlying components and estimates which components are shared across the signal. Here, it is assumed that the online reference influences all channels equally. Based on this reasoning, one can inspect the resulting mixing matrix of the independent components and remove the component with the most globally shared weights across all channels, before back-projecting the remaining data to obtain a ‘clean’ version of the original data. Whilst neighbour driven re-referencing methods (such as bipolar) are widely used in the field to maximise local signals (Li, et al., 2018; Liu, et al., 2021), it has been demonstrated that ICA might outperform bipolar re-referencing in both sensitivity and specificity (Michelmann, et al., 2018, i.e. isolating the signal from local sources and discarding activity from global sources), especially in signals of varying locality and with varying degrees of noise.

### 2.2. Data acquisition and pre-processing

#### 2.2.1 Associative memory paradigm

Data was acquired from 16 epileptic patients across two hospitals in Birmingham, UK, and Erlangen, Germany. Patients implanted with Behnke-Fried depth electrodes performed an associative memory paradigm (see Figure 3 for description) using a laptop. See Roux et al. (2022) and Kolibius et al. (2023) for details of the experimental procedure.

**Figure 3:**
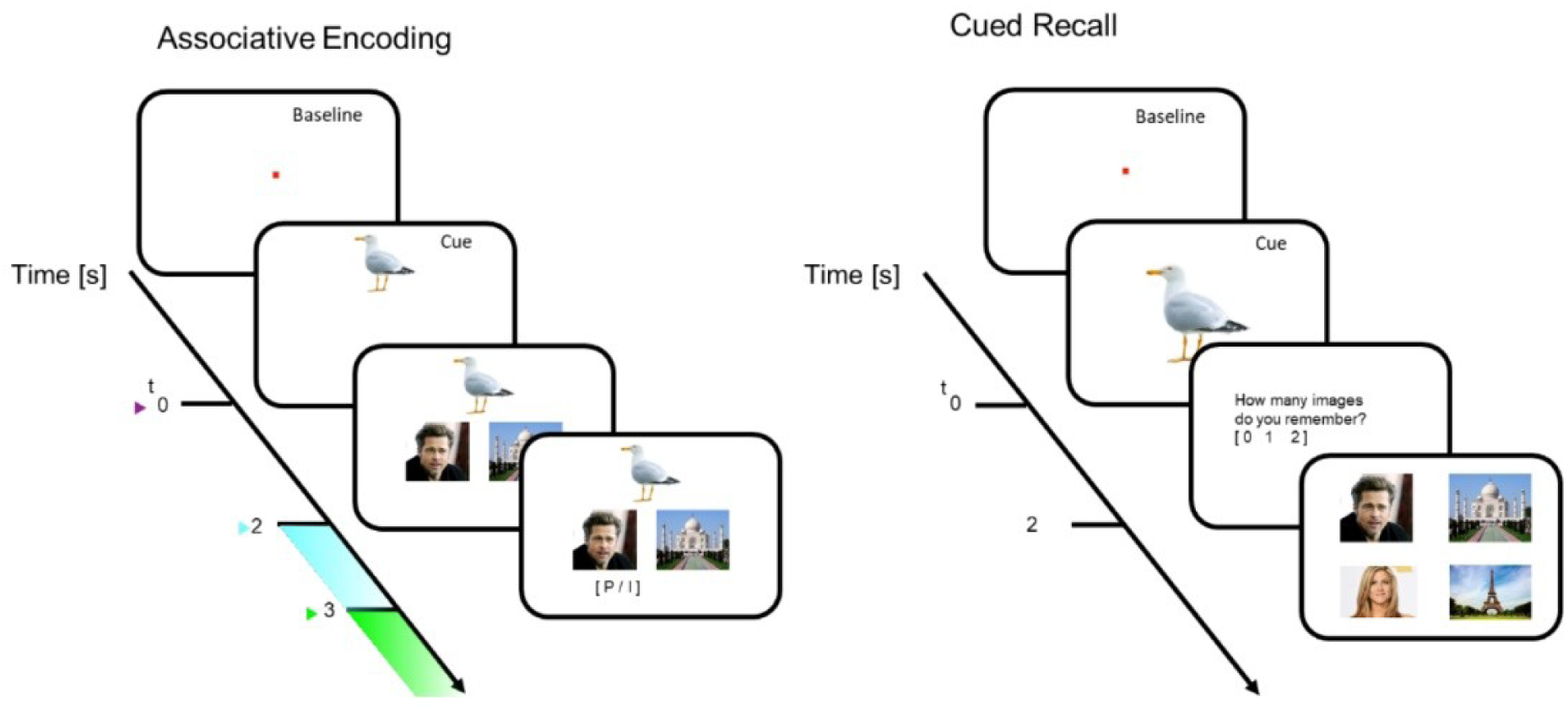
Associative memory paradigm. During an encoding phase, trials of paired images consisting of a random combination of faces and/or places were associated with a random cue (always an animal image). Each cue was later presented on its own during a retrieval phase, and the subject was asked how many images they remembered. After selecting from either zero, one, or two remembered images, the subject was then presented with a combination of four faces and/or places – two of which were the correct associated images and two of which were random. The subject was asked to identify the associated images (regardless of answer to prior question), where a trial was counted as a hit only if both associated images were recalled successfully.

#### 2.2.2 Data pre-processing

Data was originally referenced using either a bipolar or a white matter or subdural monopolar method. Recordings were resampled to 1,000 Hz before filtering and re-referencing. Line noise (N.B. in Europe line noise is at 50Hz, not 60Hz) was corrected separately for each channel and trial. A multitaper spectral estimate spanning 40–60 Hz was used to identify the dominant line-noise frequency within 47–53 Hz. When line noise was detected, its non-stationary periodic waveform was estimated using a 10 s moving-window kernel and subtracted from the signal. Detection and subtraction were repeated until the line-noise criterion was no longer met, with a maximum of 25 passes. The corrected signals were then low-pass filtered at 300 Hz and high-pass filtered at 0.2 Hz. The re-referencing methods described above were applied after these preprocessing steps.

#### 2.2.3 Artefact detection

An artefact was here considered to be an undesirable sharp transient in the electrical signal, caused by either electrical interference or interictal epileptiform discharges (IEDs) produced by pathological tissue. We considered that a trial contained an artefact if it had a variance, calculated as the root mean square (RMS) or Z-score normalisation of the LFP, above a threshold of 4. All trials with such artefacts were discarded. Any channels that had either more than 60% trials removed or five or fewer trials remaining were also removed from further analyses.

### 2.3. Data analyses

#### 2.3.1. Time frequency analyses of power

Time-frequency power was estimated from −0.5 to 5.5 s relative to cue onset, separately for low frequencies (1–15 Hz in 0.5-Hz steps; six-cycle wavelets) and high frequencies (30–100 Hz in 1-Hz steps; 12-cycle wavelets). Aperiodic components were estimated separately for each channel and analysis condition using SpecParam 2.0.0rc6. The low-frequency spectrum was parameterized over 2–15 Hz using a fixed aperiodic model, whereas the high-frequency spectrum was parameterized over 30–100 Hz using a knee model. Corrected power was calculated by subtracting the fitted aperiodic component from the observed spectrum in linear-power units. The fitted background was also retained as an outcome in its own right: its offset (the level of the fitted background) and exponent over 2–15 Hz, and its offset and effective tilt over 30–100 Hz, where the effective tilt is the slope of the fitted background across the fitted range in log–log coordinates; a larger tilt is a steeper decline of the background with frequency, and it equals the exponent when the fitted knee is zero.

##### Inter-trial phase consistency (ITPC)

Inter-trial phase consistency (ITPC) was calculated separately for each recording channel from non-artefactual trials. Channels with more than 60% rejected trials or five or fewer remaining trials were excluded. The retained signals were decomposed using FieldTrip’s wavelet method (seven-cycle wavelets) at 2-20 Hz in 0.5-Hz steps and from −0.5 to 5.5 s in 50-ms steps, using 16-s padding. Instantaneous phase was obtained from the complex Fourier coefficients, and ITPC was calculated as the magnitude of the mean unit-length phase vector across trials at each frequency-time point: ITPC(f,t) = |N^-1 sum_k exp(i phi_k(f,t))|. At every frequency-time point, phase non-uniformity across trials was assessed using Rayleigh’s test (circ_rtest, CircStat2012a). Within each channel, the complete frequency-by-time vector of p-values was adjusted with the Benjamini–Hochberg procedure. A channel was retained as ITPC-responsive when at least one adjusted p-value was ≤ 0.05. Analyses of hippocampal channels applied this criterion, whereas the supplementary all-channel analyses did not. ITPC was calculated from re-referenced LFP phase and was independent of SpecParam correction.

#### 2.3.2. Spike field coupling

For each channel, spike clusters were identified using the WaveClus toolbox (Chaure et al., 2018). Candidate LFP centre frequencies were obtained by applying MATLAB findpeaks to the all-trial 1-15 Hz residual spectrum after SpecParam aperiodic correction. For each detected centre frequency f, the original re-referenced time-domain LFP was band-pass filtered over f +/-1.25*sqrt(f) Hz and converted to analytic phase with FieldTrip’s Hilbert option. LFP phases were sampled at spike times and pairwise phase consistency (PPC; Vinck et al., 2010) was calculated. PPC was calculated separately for all trials, encoding hits and misses, and retrieval hits and misses, with at least five trials required for each subset. The same all-trial centre-frequency set was used for hit and miss calculations. Pairings were classified as same microwire, same Behnke-Fried shaft excluding same-microwire pairs, or different shaft. Main memory analyses used hippocampal LFP channels that met the ITPC-responsiveness criterion, whereas Supplementary Figures S1, S2, S4 and S5 used all eligible medial temporal channels.

#### 2.3.3. Statistical analysis

Statistical inference was tied to an explicitly defined observational unit. For the retention analyses in Figures 4 and 8, session was the repeated-measures unit and re-referencing method was the within-session factor (seven levels for macro contacts and six for microwires). All other analyses were fitted as linear mixed-effects models on individual observations. The unit of analysis was the individual channel for power and for the aperiodic parameters, and the individual spike–LFP pair for coupling (i.e., no averaging within participants). Because each re-referencing method applies one shared transformation to all channels of a shaft (AV, MD and GS each remove a signal common to the shaft, ICA removes one shared component, and WM subtracts one contact) channels from the same participant are not independent with respect to the method factor, and treating them as independent replicates would overstate differences between methods. Random intercepts for participant, session, channel or pair and participant × method, with participant × band for power and coupling and the LFP channel and spike cluster for coupling, therefore allowed errors to be correlated within these groupings (Supplementary Table S8). Where an estimated variance reached zero the term was removed under a prespecified rule, with every removal recorded in Supplementary Table S6.

**Figure 4:**
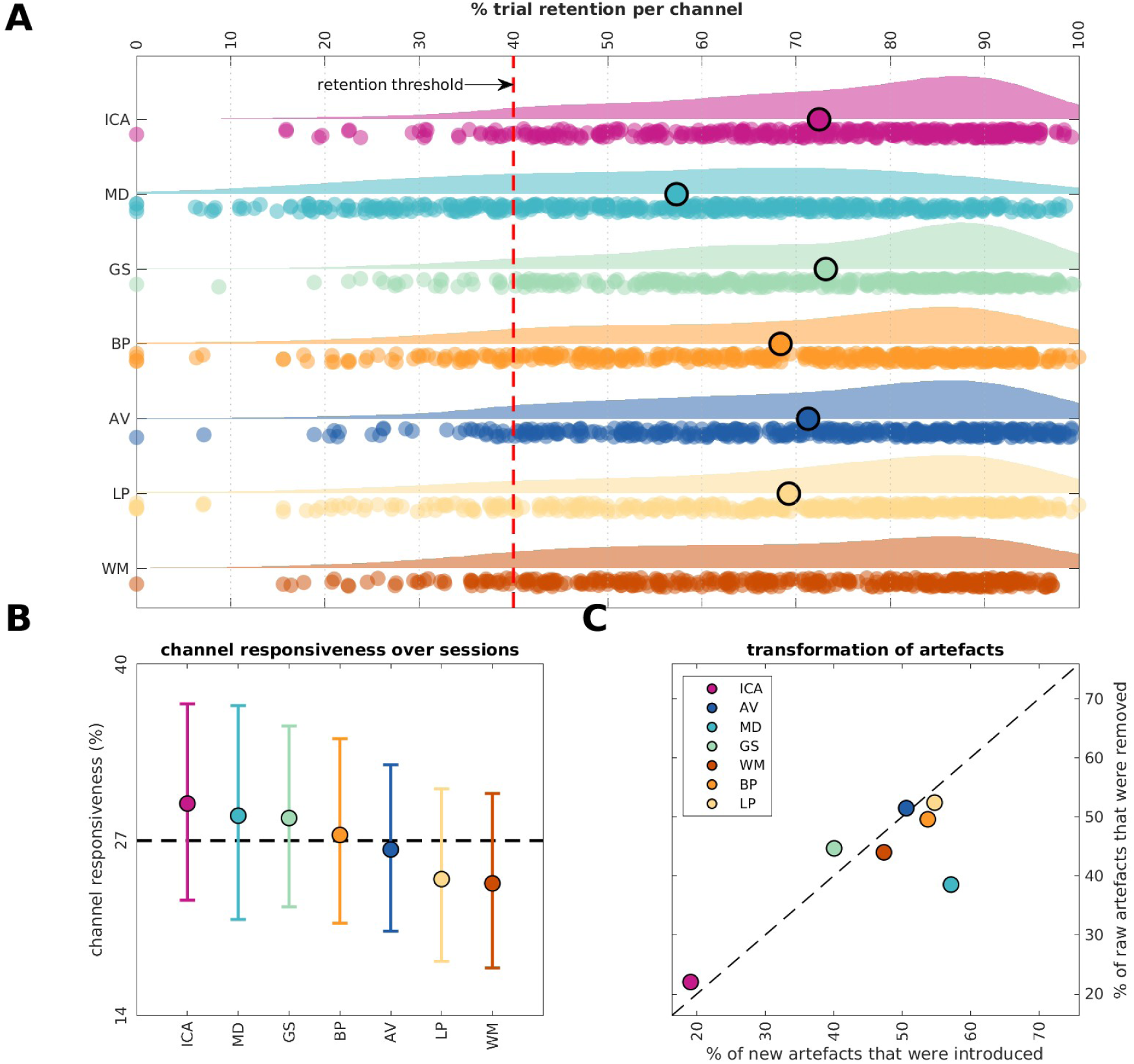
Automatic artefact rejection for macro-contact data from 54 sessions in 16 participants. Trials were indexed as artefactual when the RMS-based variance or Z-score criterion exceeded 4. Channels were excluded from subsequent analyses if more than 60% of trials were removed or if five or fewer trials remained. A: Channel-level trial-retention distributions for independent component analysis (ICA), average (AV), median (MD), Gram-Schmidt (GS), white matter (WM), bipolar (BP), and Laplacian (LP); the red dashed line marks 40% retention. The repeated-measures analysis used session-level means from the 53 sessions complete across all methods. B: Session-level percentages of eligible channels meeting the 2–20 Hz ITPC-responsiveness criterion. Points and central 90% bootstrap intervals summarize all available sessions for each method. The Benjamini–Hochberg-adjusted Greenhouse–Geisser repeated-measures omnibus test used the 45 sessions complete across all methods (Results). C: For each method, the mean percentage of raw-data artefacts absent after re-referencing is plotted against the mean percentage of post-re-referencing artefacts absent from the raw data. The dashed identity line denotes equal displayed percentages, not zero net artefact change, because the axes use different denominators. Inferential net artefact change was calculated on the common trial denominator as described in Methods

Fixed effects were re-referencing method, frequency band and their interaction for power and coupling. The aperiodic parameters have one value per fit and were modelled with method alone. Models were fitted by restricted maximum likelihood with effects coding, and tests used Satterthwaite degrees of freedom. The primary test for each power and coupling panel was the method × band interaction, and for each aperiodic panel the method effect. Planned within-band pairwise comparisons between methods, and, for the memory panels, tests of each method’s memory effect against zero, were derived from the same fitted model. Every effect that survived adjustment, including each supported omnibus test, was re-estimated with each participant removed in turn, and the results that depend on a single participant are identified where they are reported (Supplementary Table S7).

Three prespecified retention outcomes were analysed separately for macro-contact and microwire data. Channel responsiveness was the percentage of eligible channels within a session that met the ITPC-responsiveness criterion of at least one 2–20 Hz frequency-time point with a Benjamini–Hochberg-adjusted Rayleigh p-value ≤ 0.05. Trial retention was the percentage of non-artefactual trials retained for each channel. Channels with five or fewer remaining trials were assigned 0% retention for this outcome, and values were averaged across available channels within a session. Net artefact change was calculated for each channel as 100 × (number of post-re-referencing artefact flags - number of raw-data artefact flags) / number of trials and then averaged across available channels within a session. Positive values indicate a net increase in flagged trials and negative values a net decrease after re-referencing. All-missing channel-method combinations were omitted within session without imputation. Each outcome was tested with a one-way repeated-measures ANOVA using sessions with finite values for every method. Greenhouse–Geisser-adjusted degrees of freedom and p-values were used to avoid relying on the sphericity assumption. The three omnibus Greenhouse–Geisser p-values were Benjamini–Hochberg-adjusted separately within the macro-contact and microwire analyses. When the adjusted omnibus p-value was ≤ 0.05, all method pairs were compared with two-sided paired-samples t-tests, with Benjamini–Hochberg adjustment across the 21 macro-contact or 15 microwire comparisons within that outcome.

For power memory effects, hit and miss trials were compared separately within each channel and frequency using a two-sided, equal-variance independent-samples t-test. The resulting t statistic was retained as the channel-wise memory-effect estimate. Low-frequency power summaries used 2.5–4, 4–8 and 8–12 Hz bands.

For coupling memory effects, PPC was calculated separately for hits and misses, and the response of each spike–LFP pair was hit PPC minus miss PPC. Coupling used the same 2.5–4, 4–8 and 8–12 Hz bands as power. Differences between methods, and the direction of each method’s memory effect, were tested in the mixed model described above.

Multiple-comparison correction depended on the analysis stage. The retention analyses used the Benjamini–Hochberg families described above. For ITPC channel selection, Rayleigh-test p-values were adjusted across the full frequency-by-time grid separately within each channel using Benjamini–Hochberg FDR. No additional correction was applied across channels or sessions. Multiple comparisons were controlled with the Benjamini–Hochberg procedure within four prespecified families: the eighteen method × band interaction tests; the twenty-four aperiodic method tests; within each panel, the planned within-band pairwise comparisons between methods; and, for the memory panels, the tests of each method’s memory effect against zero. Every test is listed in Supplementary Tables S2–S4. An adjusted p ≤ 0.05 is reported as an effect and 0.05 < p ≤ 0.10 as a trend, and the two are marked distinctly in the figures. The sensitivity analysis used identical families. Statistical significance was evaluated at p ≤ 0.05.

Intervals accompanying model estimates are pointwise 95% confidence intervals from the fitted model; the descriptive spectra in Figures 5, 6, 9 and 10 instead show the 2.5th to 97.5th percentiles of 2,000 bootstrap resamples across channels. Mixed models and the retention analysis were fitted in MATLAB (R2023a).

**Figure 5:**
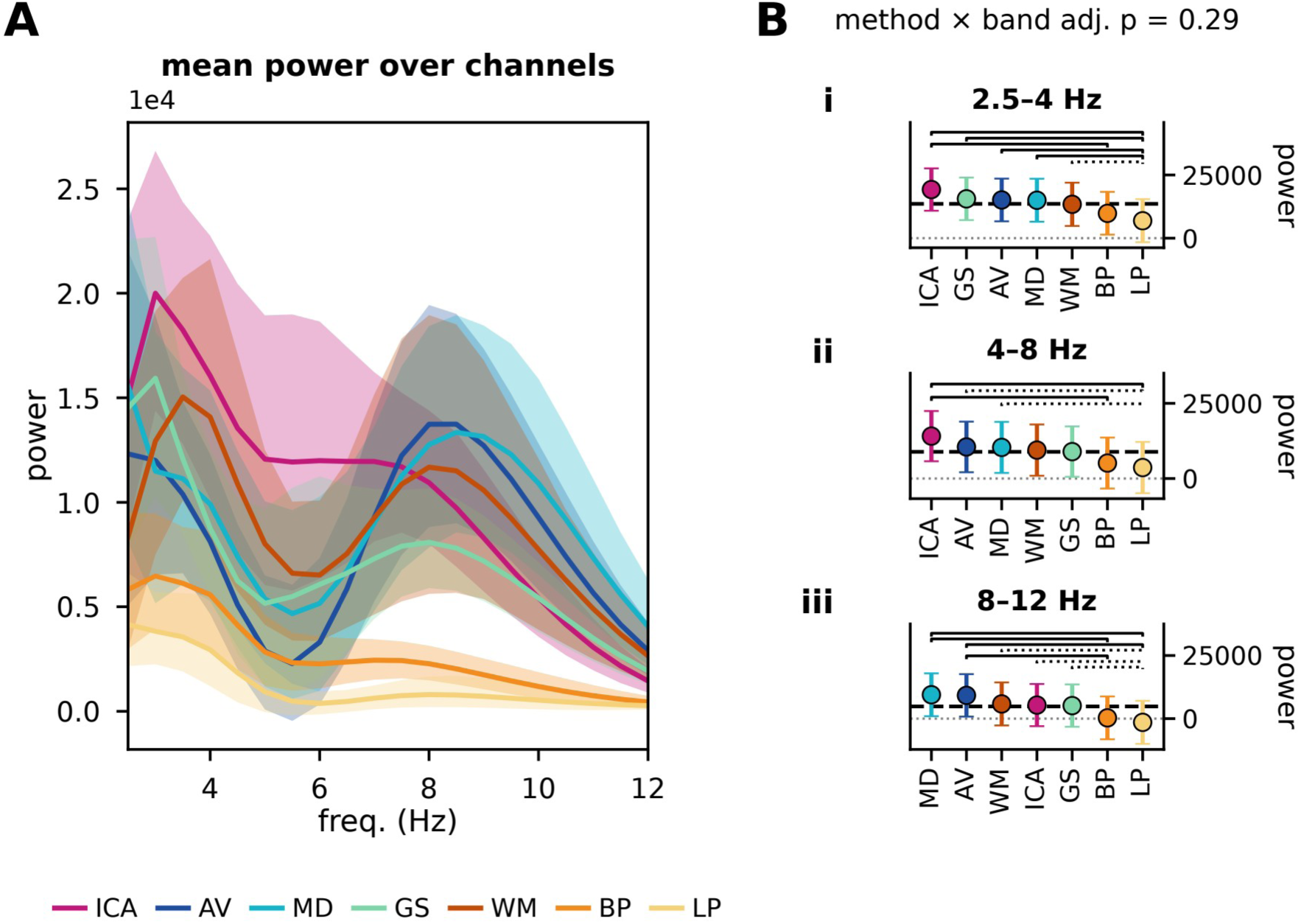
Corrected power summaries for ITPC-responsive hippocampal macro-contact channels across seven re-referencing methods: independent component analysis (ICA), average (AV), median (MD), Gram-Schmidt (GS), white matter (WM), bipolar (BP), and Laplacian (LP). A: mean corrected power per method over 2.5–12 Hz, in linear-power units; the shaded band is the 2.5th to 97.5th percentile of 2,000 bootstrap resamples across channels. B: corrected power in 2.5–4, 4–8 and 8–12 Hz bands, as model-estimated means with pointwise 95% confidence intervals, methods rank-ordered by estimate; brackets mark pairwise comparisons that pass Benjamini–Hochberg within the panel’s family (solid, adjusted p ≤ 0.05; dotted, 0.05 < adjusted p ≤ 0.10), and the method × band interaction is given above the band panels. The unit of analysis is the individual channel: ICA 89, AV 71, MD 56, GS 84, WM 69, BP 78 and LP 70 channels from 15 participants. The band at 2 Hz is the lower boundary of the aperiodic fitting range and is excluded throughout (Methods).

**Figure 6:**
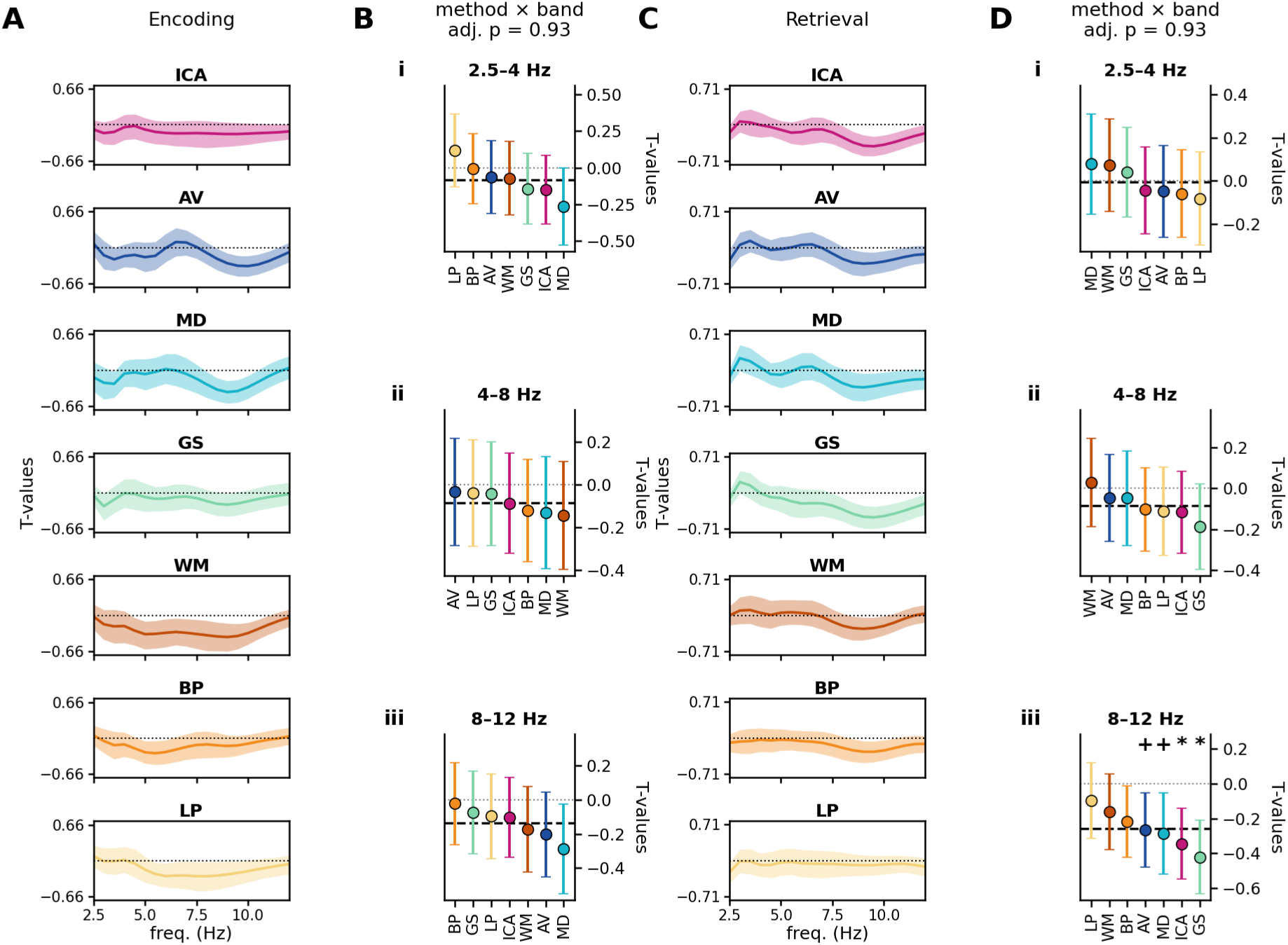
Power memory effects for ITPC-responsive hippocampal macro-contact channels (Methods), across seven re-referencing methods. Encoding is shown in A–B and retrieval in C–D. Within each channel and frequency, hits and misses were compared with a two-sided equal-variance independent-samples t-test; the resulting t statistic is the channel-wise memory-effect estimate. A and C show the mean of those estimates per method over 2.5–12 Hz, with the 2.5th to 97.5th percentile of 2,000 bootstrap resamples across channels; they are descriptive. B and D show the estimates in 2.5–4, 4–8 and 8–12 Hz bands as model-estimated means with pointwise 95% confidence intervals, methods rank-ordered by estimate; brackets mark pairwise comparisons, and a star (adjusted p ≤ 0.05) or a plus sign (trend) marks a method whose memory effect differs from zero, both after Benjamini–Hochberg within the panel’s two families; the method × band interaction is given above B and D. The unit of analysis is the individual channel: 130 channels from 14 participants at encoding and 129 from 15 at retrieval.

**Figure 7:**
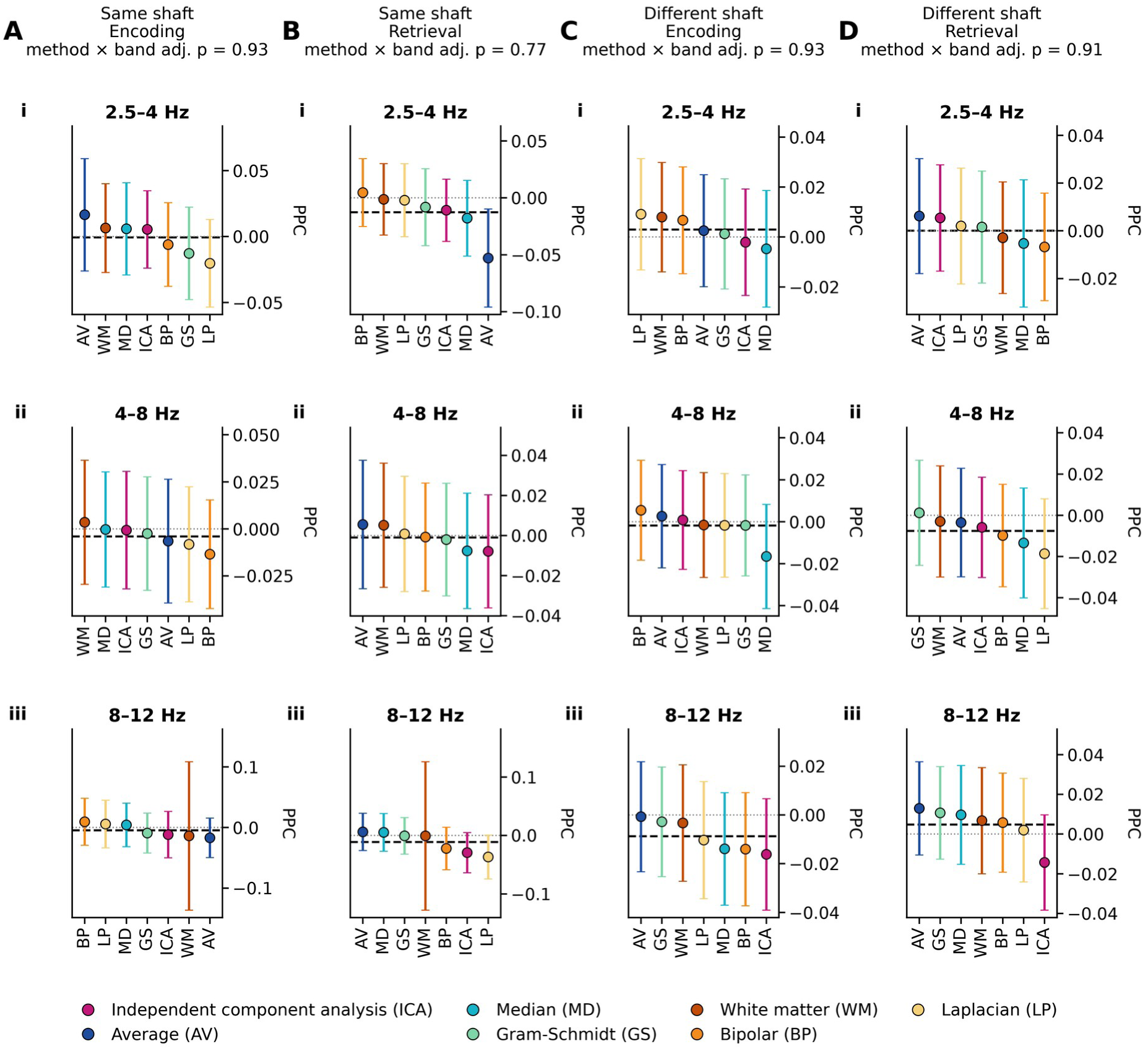
Spike–LFP coupling memory effects for ITPC-responsive hippocampal macro-contact LFP channels across seven re-referencing methods. Candidate centre frequencies were detected from each channel’s all-trial corrected spectrum; pairwise phase consistency (PPC) for hits and misses was then computed from the re-referenced time-domain LFP at the same centre frequencies, and the memory effect is hit minus miss. Every finite PPC estimate was retained. Panels show same-shaft pairings (A: encoding; B: retrieval) and different-shaft pairings (C: encoding; D: retrieval) in 2.5–4, 4–8 and 8–12 Hz bands, as model-estimated means with pointwise 95% confidence intervals, methods rank-ordered by estimate. Each column header gives the method × band interaction. No pairwise comparison and no memory effect passed Benjamini–Hochberg within the panel’s families. The unit of analysis is the individual spike–LFP pair: 368 to 1,384 pairs per panel from 11 to 13 participants.

Corrected power and spike–LFP coupling were summarised in three bands: 2.5–4, 4–8 and 8–12 Hz. The lowest band starts at 2.5 Hz because 2 Hz is the lower boundary of the aperiodic fit, where corrected power is systematically underestimated. Spike–LFP pairs whose coupling was estimated only at frequencies below 2.5 Hz therefore contribute no lowest-band observation (33 of 93,804 observations in the main analysis and 1,023 of 764,829 in the supplementary analysis). Five of 97,568 aperiodic fits were excluded because the fitted background was undefined or numerically unstable; the sensitivity analysis additionally excluded 948 fits with a fitting error above 0.15 or an RZ below 0.85.

## 3 Results

Results are split into two sections, based on the type of electrode (macro electrodes or micro-wire electrodes). Each section comparatively analyses the effect of up to seven different re-referencing methods on SEEG data. The data were analysed for differences between re-referencing schemes with respect to artefact detection, oscillatory power, and modulation of oscillatory power by behaviour.

### 3.1 Macro contacts

#### 3.1.1 Artefact detection

Artefacts occur in electrophysiological data for a number of reasons. One such source might be electrical interference near to the recording site, which can be more stationary in the form of 50 Hz line noise or more non-stationary for other forms of electrical bursts such as induced by muscle activity. Another source might be what are known as interictal epileptic discharges (IEDs), internally generated electrophysiological bursts that are caused by pathological tissue. Trials that exhibit these forms of interference are commonly removed from the analyses pipeline as they disrupt analyses of the signal. Here, an automatic artefact rejection algorithm was used to identify and remove such trials (Figure 4A), based on analysing the variance of a given trial in relation to other trials.

As re-referencing transforms the signal, it can change which trials cross the automatic artefact-rejection threshold. We therefore quantified both the proportion of trials retained after artefact rejection and how the set of artefact flags changed relative to the raw data.

Re-referencing method affected session-level trial retention (Figure 4A; 53 complete sessions; Greenhouse–Geisser-corrected repeated-measures ANOVA, F(3.04, 158.33) = 58.89, adjusted p < 0.001). Mean session-level retention was 68.3% for GS, 67.8% for ICA, 66.8% for AV, 65.5% for WM, 62.5% for BP, 62.2% for LP, and 51.6% for MD. Adjusted paired comparisons showed lower retention for MD than for every other method (all p < 0.001). Among the remaining methods, retention was higher for GS than AV, BP, LP and WM; for ICA than BP, LP and WM; for AV than BP and LP; and for WM than BP and LP (all adjusted p ≤ 0.0259).

Figure 4C describes this transformation from two directions. For MD, an average of 57.1% of artefact flags detected after re-referencing were not present in the raw data, whereas 38.5% of raw-data artefact flags were no longer detected after re-referencing. Because these two percentages use different denominators—post-re-referencing artefact flags and raw-data artefact flags, respectively—they cannot be subtracted directly. Inferential comparisons therefore used the common-denominator net artefact-change score.

The common-denominator net artefact-change score also differed among methods (Figure 4C; 53 complete sessions; F(3.04, 157.94) = 58.04, adjusted p < 0.001). Mean changes, in percentage points of trials, were −0.9 for GS, −0.4 for ICA, 0.6 for AV, 2.2 for WM, 4.9 for BP, 5.1 for LP, and 15.7 for MD. MD produced a larger net increase than every other method (all adjusted p < 0.001). Additional significant comparisons showed lower net change for AV than BP and LP; for GS than AV, BP, LP, and WM; for ICA than BP, LP, and WM; and for WM than BP (all adjusted p ≤ 0.0073). Because this score and trial retention are derived from the same post-re-referencing artefact flags, the similar pattern does not constitute an independent confirmatory result. Taken together, the lower trial retention and larger positive net artefact change for MD show that the automatic procedure flagged more trials after MD re-referencing.

In contrast, channel responsiveness did not differ reliably among the seven macro-contact methods (Figure 4B). The all-available descriptive summaries shown in the panel ranged from 23.8% for WM to 29.7% for ICA. The repeated-measures comparison used the 45 sessions with values for every method and was non-significant (F(3.54, 155.59) = 1.53, Greenhouse–Geisser-corrected p = 0.2034). Thus, the data do not establish that ICA or any other method retained a greater proportion of responsive macro-contact channels.

#### 3.1.2 Power

Figure 5 summarizes low-frequency oscillatory power after SpecParam aperiodic correction. The plotted corrected quantity is observed linear power minus the fitted aperiodic component in linear-power units. Methods were compared at the band level in the mixed model, in which method and band did not interact (F(12, 1286) = 1.89, adjusted p = 0.29). The frequency-resolved curves are descriptive.

Figure 5A indicates that in general these lower frequencies are split into three components (see Figure 5B). The first two could be considered part of the theta rhythm that originates in the MTL (typically 2-8Hz, here exhibited by two 2.5–4 Hz & 4–8 Hz components) (von Stein & Sarnthein, 2000; Buzsáki & Draguhn, 2004; Lega et al., 2012; Goyal et al., 2020; Roux et al., 2022). These components are most reduced by the highly localised Laplacian method, which has previously been shown to isolate local task-related cortical activity (Li et al., 2018, Liu et al., 2021). Its corrected power was the lowest in all three bands, and at 2.5–4 Hz it was below ICA, AV, MD and GS (all adjusted p ≤ 0.027; the AV comparison depends on one participant; WM, trend, adjusted p = 0.086).

The third could be considered part of the alpha rhythm (here exhibited by an 8-12Hz component) which is typically dominant in neocortical areas (Griffiths et al., 2019). Interestingly, alpha oscillations are strongest in the more global re-referencing methods, such as the mean [AV] or median [MD] average, whereas it is most reduced in the local re-referencing schemes such as LP and BP (Fig. 5B, 8–12 Hz; AV and MD each exceeded BP and LP, all adjusted p ≤ 0.011). ICA, GS and WM lay between these extremes: none differed from AV or MD (all adjusted p ≥ 0.21), each exceeded LP only as a trend (adjusted p = 0.058 –0.066), and none differed from BP. It is conceivable that AV and MD introduced this signal from neocortical regions. This would also be in line with previous studies that found global re-referencing (i.e. AV and MD) to similarly contaminate local signals with other more global signals (Li et al., 2018).

The fitted background itself differed between methods (Supplementary Figure S3). Its offset over 2–15 Hz was higher under ICA, AV, MD and GS than under BP and LP (F(6, 53) = 9.59, adjusted p < 0.001; all pairwise adjusted p ≤ 0.010), with WM intermediate — above LP (adjusted p = 0.007) but not above BP. The exponent over the same range, and both parameters over 30–100 Hz, showed no method effect (all adjusted p ≥ 0.114). Of their planned comparisons, two passed: the effective tilt over 30–100 Hz was higher under AV than under ICA and under LP (both adjusted p = 0.038). The offset ranks the methods in the same order as the corrected power in the two theta bands — ICA highest, BP and LP lowest, with AV, MD and GS indistinguishable from one another in both (all adjusted p ≥ 0.554 for the background and ≥ 0.70 for theta power). Re-referencing therefore does not trade the background against the oscillatory power above it; in the theta bands it moves both together. This is consistent with the account of these methods given above: the more of the signal shared across neighbouring contacts a method subtracts, the lower the background it leaves and the less power remains in the bands above it.

#### 3.1.3 Memory effects

Next, we analyse how the quality of oscillatory signals (using both power and spike field coherence) is modulated by behaviour. Here, a cued recall paradigm was conducted that required patients to form and retrieve associative memories between three images (see Roux et al. 2022; Kolibius et al., 2023, and Figure 3 for design). Of main interest were difference dependent on memory success, that is whether the memory was complete (i.e. both images correctly recalled), or incomplete (one or both images incorrect). Complete memories are called ‘hits’, whereas incomplete memories are called ‘misses’. We analysed difference in power or spike field coherence between hit and miss trials, for both the encoding and retrieval phases of the paradigm. Therefore, it is important to consider whether or not re-referencing has any bearing on such memory effects.

##### 3.1.3.1 Memory effects in power

Overall, the band-level comparisons identified a negative alpha (8–12 Hz) memory effect at retrieval, indicating higher power for misses than hits (Herweg et al., 2020). Re-referencing method and frequency band did not interact at either memory stage (encoding F(12, 1021) = 0.81, retrieval F(12, 1023) = 0.80, both adjusted p = 0.93), and no method differed from another in any band or at either stage. Against zero, the retrieval alpha effect was supported under ICA (−0.34, adjusted p = 0.010) and GS (−0.42, adjusted p = 0.003), with AV and MD as trends (both adjusted p = 0.082); no effect was supported at 2.5–4 or 4–8 Hz, or at encoding. The memory related alpha power decreases (Staresina et al., 2016) do not establish whether the underlying source is local or spatially distributed. The fitted background gave the same picture (Supplementary Figure S3): no method differed from another in any parameter at either stage (one trend, ICA below LP for the 30–100 Hz offset at encoding, adjusted p = 0.095). Against zero, only the 2–15 Hz exponent at retrieval was supported: the fitted background was steeper for hits than for misses under GS, BP and LP (each adjusted p = 0.049, each dependent on one participant), with ICA and AV as trends (both adjusted p = 0.052).

##### 3.1.3.2 Memory effects in spike field coupling

Spike–LFP coupling gave no evidence that re-referencing alters memory effects on macro contacts. Method and band did not interact for same-shaft or different-shaft coupling at either stage (all four adjusted p ≥ 0.77), no method differed from another in any band, and no method’s memory effect differed from zero.

### 3.2 Micro-wires

#### 3.2.1 Artefact detection

Re-referencing method also affected session-level trial retention for microwires (Figure 8A; 54 sessions; F(3.64, 193.02) = 76.07, adjusted p < 0.001). Mean session-level retention was 61.2% for ICA, 59.0% for AV, 58.1% for LP, 58.0% for BP, 57.4% for GS, and 48.3% for MD. Adjusted paired comparisons showed higher retention for ICA than for every other method (all p ≤ 0.0045), lower retention for MD than for every other method (all p < 0.001), and higher retention for AV than GS (p = 0.0103).

Net artefact change differed correspondingly among microwire methods (Figure 8C; F(3.65, 193.52) = 76.34, adjusted p < 0.001). Mean changes, in percentage points of trials, were 4.1 for ICA, 6.4 for AV, 7.2 for LP, 7.4 for BP, 7.9 for GS, and 17.0 for MD. ICA showed a smaller net increase than every other method (all adjusted p ≤ 0.0044), MD showed a larger net increase than every other method (all p < 0.001), and GS showed a larger increase than AV (p = 0.0107). As in the macro-contact analysis, this outcome is mathematically linked to trial retention and is not an independent confirmation.

Microwire channel responsiveness also differed among methods (Figure 8B; 54 sessions; F(3.34, 177.20) = 5.15, Greenhouse–Geisser-corrected p = 0.0013, adjusted p = 0.0013). Mean responsiveness was 21.7% for ICA and 14.2–15.9% for the other methods. Adjusted paired comparisons indicated greater responsiveness for ICA than every other method (all p ≤ 0.0215). No comparison among the remaining five methods was significant. ICA therefore retained a greater proportion of responsive microwire channels than any alternative, whereas the same comparison on macro contacts was non-significant.

**Figure 8:**
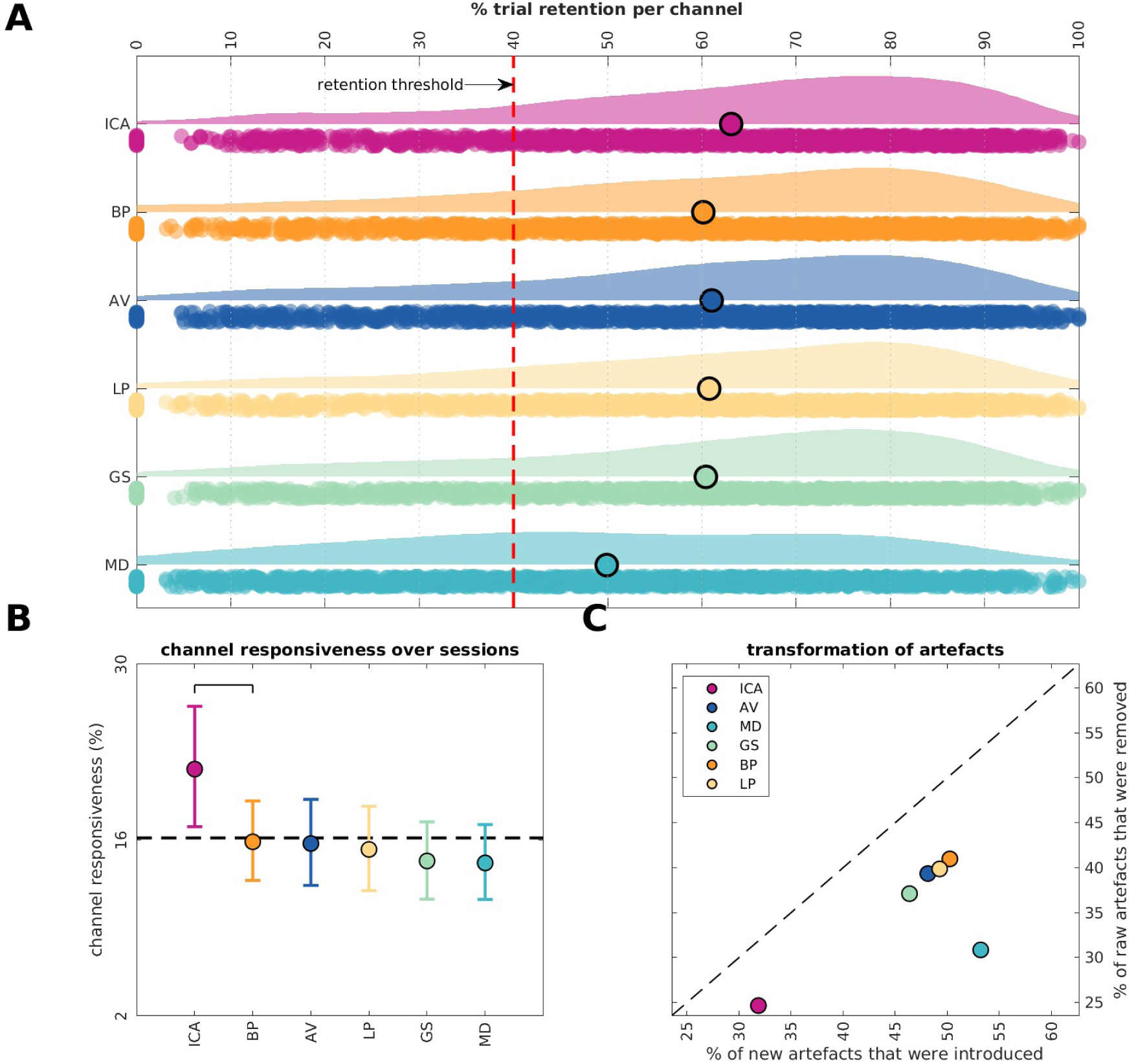
Automatic artefact rejection for microwire data from 54 sessions in 16 participants. Artefact and channel-exclusion criteria were as in Figure 4. A: Channel-level trial-retention distributions for independent component analysis (ICA), average (AV), median (MD), Gram-Schmidt (GS), bipolar (BP), and Laplacian (LP); the red dashed line marks 40% retention. The repeated-measures analysis used session-level means from all 54 sessions. B: Session-level percentages of eligible channels meeting the 2–20 Hz ITPC-responsiveness criterion, with central 90% bootstrap intervals. The Benjamini–Hochberg-adjusted Greenhouse–Geisser repeated-measures omnibus test used all 54 sessions (Results). In B, methods are rank-ordered, and a horizontal bracket runs from a method to the nearest method it differs from after adjustment; it therefore also differs from every method beyond that one. C: Mean percentages of removed and introduced artefacts, defined as in Figure 4. The dashed identity line denotes equal displayed percentages; the session-level net artefact-change test used a common trial denominator.

#### 3.2.2 Power

Continuing this pattern, ICA yielded higher corrected power than AV, GS, LP and MD at 2.5 –4 Hz and than AV, GS and LP at 4–8 Hz (all adjusted p ≤ 0.030; against BP, adjusted p = 0.085 at 2.5–4 Hz and not supported at 4–8 Hz; Figure 9B). Unlike the macro contacts, method and band interacted (F(10, 2187) = 9.63, adjusted p < 0.001), so the ordering is band-specific: BP ranked second at 2.5–4 and 4–8 Hz but highest at 8–12 Hz, where ICA ranked lowest. One notable difference to the application of the same re-referencing method on the macro contact data is the relatively higher ranking of neighbour driven methods in low-frequency power. This might be since it is impossible to practically identify neighbouring micro-wires in MRI scans, and so these methods should more properly be considered as randomised monopolar methods here. This better performance might also reflect a higher degree of independence amongst micro-wires, which are highly localised and thus might share less global low frequency rhythms. AV and GS had the lowest mean corrected power in the two theta bands.

**Figure 9:**
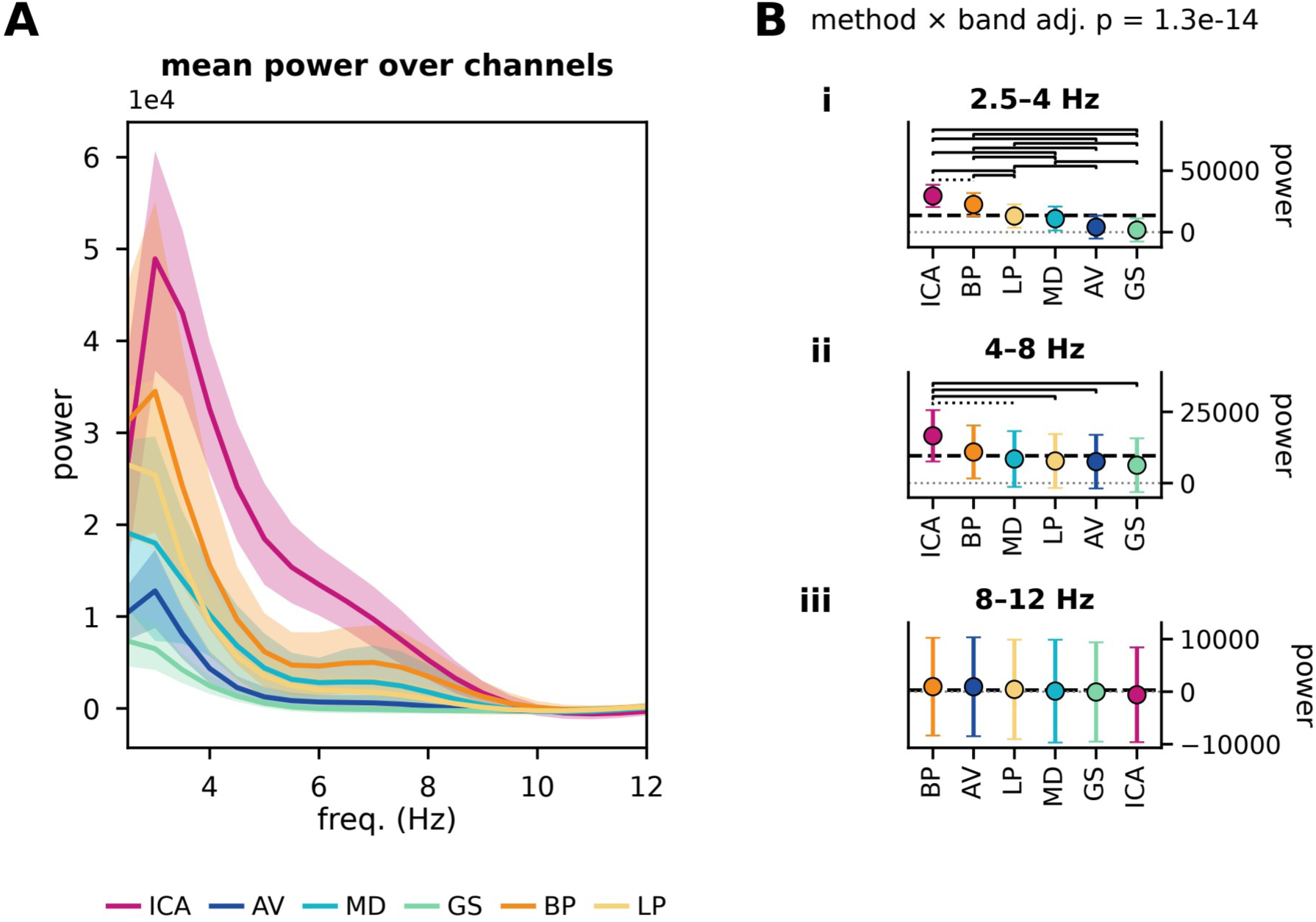
Corrected power summaries for ITPC-responsive hippocampal microwire channels across six re-referencing methods: independent component analysis (ICA), average (AV), median (MD), Gram-Schmidt (GS), bipolar (BP), and Laplacian (LP). A: mean corrected power per method over 2.5–12 Hz, in linear-power units; the shaded band is the 2.5th to 97.5th percentile of 2,000 bootstrap resamples across channels. B: corrected power in 2.5–4, 4–8 and 8–12 Hz bands, as model-estimated means with pointwise 95% confidence intervals, methods rank-ordered by estimate; brackets mark pairwise comparisons that pass Benjamini–Hochberg within the panel’s family (solid, adjusted p ≤ 0.05; dotted, 0.05 < adjusted p ≤ 0.10), and the method × band interaction is given above the band panels. Method and frequency band interacted for these channels (F(10, 2187) = 9.63, adjusted p < 0.001), so the ordering of methods differs between bands.

As for the macro contacts, the fitted background differed between methods, and here it did so in every parameter measured (Supplementary Figure S3). Its offset over 2–15 Hz was higher under ICA, BP and LP than under AV, MD and GS (F(5, 57) = 11.14, adjusted p < 0.001; all pairwise adjusted p ≤ 0.032), and its exponent over the same range was lower under GS than under ICA, AV, BP and LP, and under MD than under ICA (F(5, 57) = 6.31, adjusted p < 0.001). Over 30–100 Hz, the offset was higher under ICA than under AV, MD and GS (F(5, 40) = 3.95, adjusted p = 0.025; this method effect depends on one participant, its three pairwise comparisons do not) and the effective tilt was higher under ICA than under MD and GS, and lower under GS than under BP and LP (F(5, 44) = 4.02, adjusted p = 0.025). As on the macro contacts, the background offset ranks the methods in the same order as the corrected power at 2.5–4 Hz — here ICA, BP and LP highest and GS lowest — but that ranking is close to inverted: BP and LP, which leave the lowest background on macro contacts, leave among the highest on microwires. This is consistent with the preceding account: on a macro shaft BP or LP subtracts a genuine neighbour and removes signal the two contacts share; microwires cannot be ordered spatially, so the same operation subtracts a wire that shares little with the target and behaves closer to a monopolar reference.

#### 3.2.3 Memory effects

##### 3.2.3.1 Memory effects in power

Re-referencing method and frequency band did not interact for microwire power memory effects at either stage (encoding F(10, 1630) = 1.30, adjusted p = 0.77; retrieval F(10, 1501) = 0.59, adjusted p = 0.93; Figure 10), and no method differed from another in any band. Tested against zero, the retrieval memory effect at 4–8 Hz was negative, i.e. power was lower for hits than for misses, under ICA (−0.32, adjusted p < 0.001) and AV (−0.27, adjusted p = 0.019). No other microwire power memory effect differed from zero. The fitted background showed no method difference at either stage (Supplementary Figure S3). At encoding, the 2–15 Hz offset under GS was higher for hits than for misses (adjusted p = 0.049 against zero; dependent on one participant). The 2–15 Hz exponent under ICA was steeper for hits than for misses as a trend (adjusted p = 0.056).

**Figure 10:**
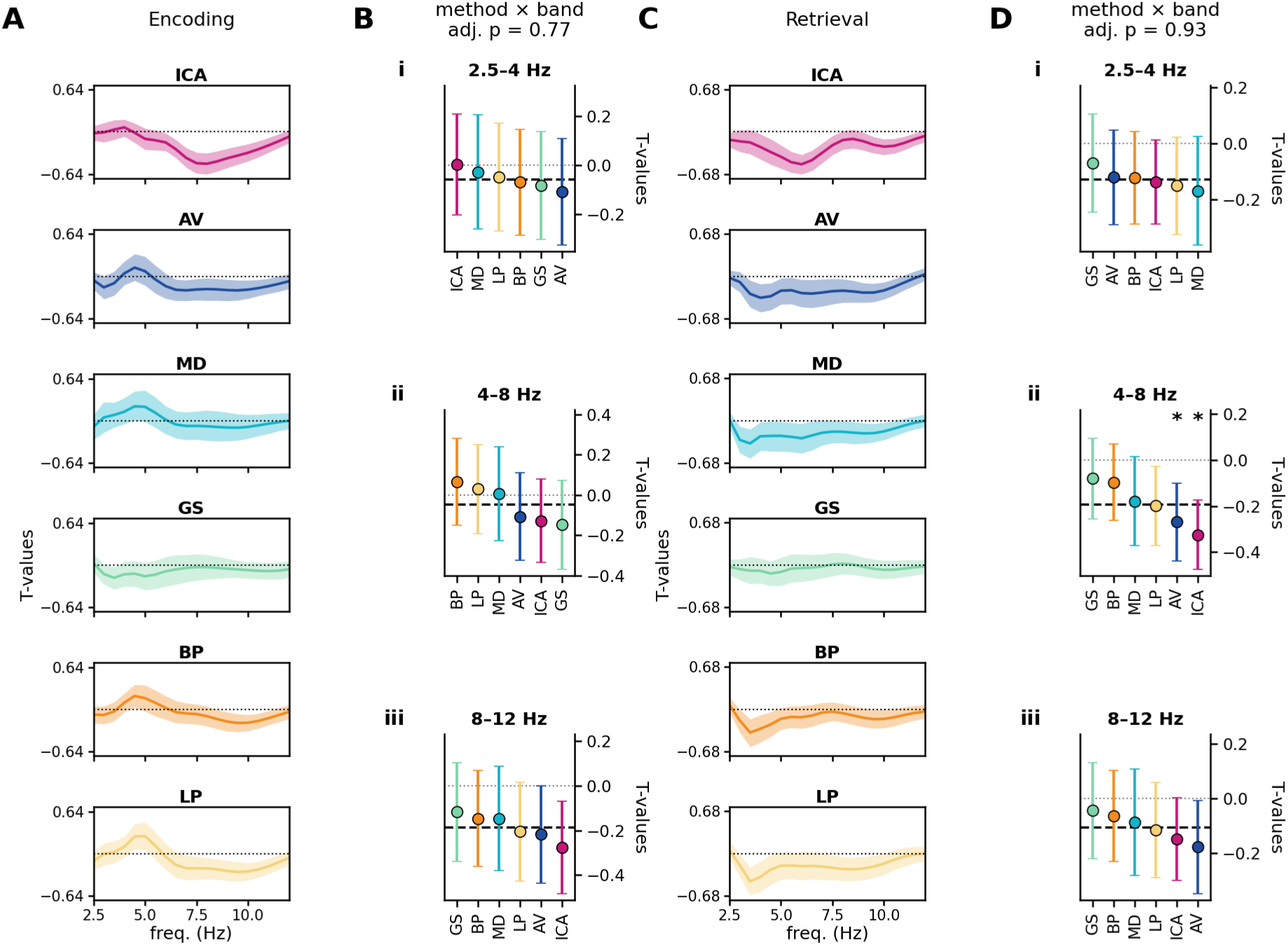
Power memory effects for ITPC-responsive hippocampal microwire channels (Methods), across six re-referencing methods. Encoding is shown in A–B and retrieval in C–D. Within each channel and frequency, hits and misses were compared with a two-sided equal-variance independent-samples t-test; the resulting t statistic is the channel-wise memory-effect estimate. A and C show the mean of those estimates per method over 2.5–12 Hz, with the 2.5th to 97.5th percentile of 2,000 bootstrap resamples across channels; they are descriptive. B and D show the estimates in 2.5–4, 4–8 and 8–12 Hz bands as model-estimated means with pointwise 95% confidence intervals, methods rank-ordered by estimate; brackets mark pairwise comparisons, and a star (adjusted p ≤ 0.05) or a plus sign (trend) marks a method whose memory effect differs from zero, both after Benjamini–Hochberg within the panel’s two families; the method × band interaction is given above B and D. The unit of analysis is the individual channel: 318 channels from 14 participants at encoding and 299 from 16 at retrieval.

**Figure 11:**
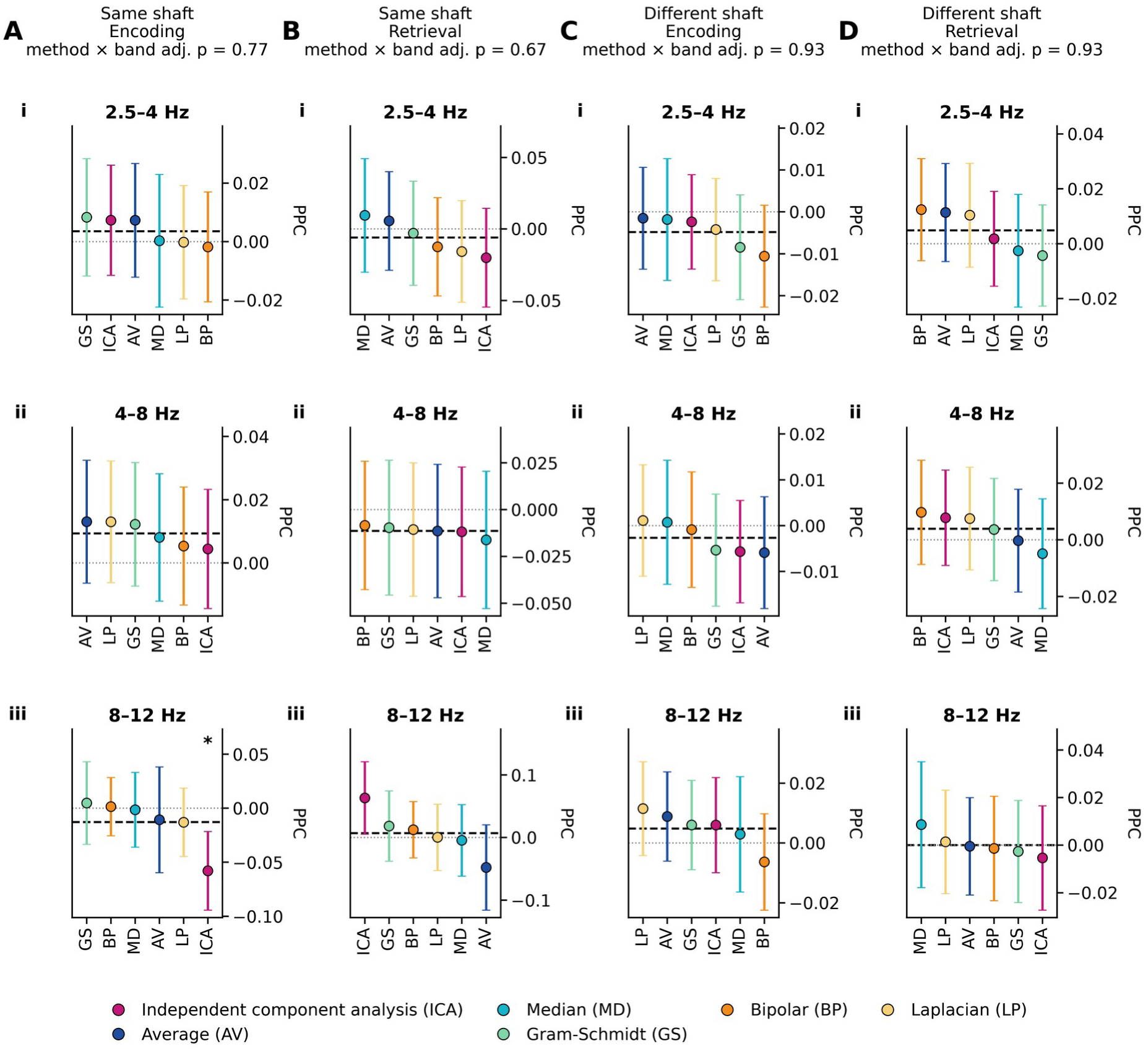
Spike–LFP coupling memory effects for ITPC-responsive hippocampal microwire LFP channels across six re-referencing methods. Candidate centre frequencies were detected from each channel’s all-trial corrected spectrum. Pairwise phase consistency (PPC) for hits and misses was then computed from the re-referenced time-domain LFP at the same centre frequencies, and the memory effect is hit minus miss. Panels show same-shaft pairings that exclude the exact same microwire (A: encoding; B: retrieval) and different-shaft pairings (C: encoding; D: retrieval) in 2.5–4, 4–8 and 8–12 Hz bands, as model-estimated means with pointwise 95% confidence intervals, methods rank-ordered by estimate. Each column header gives the method × band interaction. No pairwise comparison passed Benjamini–Hochberg within the panel’s families. A star marks the one memory effect that differs from zero after adjustment: same-shaft coupling at 8–12 Hz under ICA at encoding, from 33 pairs in five participants, which does not survive the removal of one participant. The unit of analysis is the individual spike–LFP pair: 760 to 3,032 pairs per panel from 10 to 14 participants.

##### 3.2.3.2 Memory effects in spike field coupling

Microwire coupling likewise gave no evidence that re-referencing alters memory effects: method and band did not interact in any of the four panels (all adjusted p ≥ 0.67), and no method differed from another in any band. One memory effect differed from zero: under ICA, same-shaft coupling at 8–12 Hz was reduced for hits relative to misses at encoding (−0.058, adjusted p = 0.035; 33 pairs from five participants). That estimate rests on few participants and does not survive the removal of one of them, so it is reported but not interpreted.

### 3.3 Summary of statistics

Method and band interacted for microwire power across all trials (F(10, 2187) = 9.63, adjusted p < 0.001) and in none of the other seventeen power and coupling panels: not for macro-contact power, not for same-shaft or different-shaft coupling, and at neither memory stage. Method effects on the aperiodic background were present in five of the twenty-four aperiodic tests, all of them in the all-trial data. Memory effects that differed from zero were confined to retrieval power — alpha on macro contacts, fast-theta (4–8 Hz) on microwires — to one microwire coupling estimate and to four aperiodic estimates, the last five each dependent on one participant; none of them differed between methods. In the supplementary population of all eligible medial temporal channels (Supplementary Figures S4 and S5), four further omnibus tests were supported: the method × band interactions for macro-contact power and for microwire different-shaft coupling, and the method effects on the macro-contact exponent over 2–15 Hz and effective tilt over 30–100 Hz. One difference between methods in a memory effect was also supported there: at encoding, different-shaft macro-contact coupling at 8–12 Hz was higher under GS than under BP and LP (adjusted p = 0.029). The random-effect estimates show where the clustering of channels within participants mattered (Supplementary Table S8). In the memory-effect models of power and coupling the participant term accounted for none of the variance in any power panel and for at most 7% in coupling, so the absence of method differences in the memory effects is a property of the data rather than of the model. In the all-trial models of the aperiodic background it accounted for up to 65% of the total variance for the macro-contact offset over 2–15 Hz and 76% for its exponent, and in all-trial power the participant and participant × band terms together accounted for 50% on macro contacts and 22% on microwires. These are the comparisons in which a test that treats channels as independent replicates would overstate the differences between methods.

No reversal between methods was supported. A reversal was counted only when two methods’ memory effects had opposite signs and both, together with their direct comparison, met the adjusted criterion; no pair met it, in either population, for power, coupling or the aperiodic parameters (Supplementary Table S5).

## 4 Concluding remarks

We set out to ask whether the choice of re-referencing method changes what is measured from macro contacts and microwires during an episodic memory task, and whether it changes the memory effects those measurements carry. Re-referencing transformed these recordings in three ways. It changed which trials survived automatic artefact rejection. It changed low frequency power, by amounts that differed across frequency bands in the microwire recordings. And it shifted the aperiodic background against which that power is read — in the theta bands the background and the power above it moving together rather than trading against one another, since the methods that left the highest background also left the most power above it. We found no evidence that re-referencing changed the detection of behavioural effects in the hippocampal recordings: no method differed from any other in any power or spike–LFP coupling memory effect, and in neither population did any pair of methods show a supported reversal in the direction of a memory effect. In general, each re-referencing method seems to work in accordance with its own assumptions about the underlying components of the signal, where different methods work differently to maximise local or global components. Though there does not seem to be a one size fits all method, there are some promising signs that data-driven methods are a versatile tool that should be further explored (for further reading see Michelmann et al., 2018 and Schaworonkow & Voytek, 2021). Previous comparative studies of this type have shown the benefits of using a localised re-referencing method (such as Laplacian) for the analyses of local cortical activity during motor tasks, where high spatial resolution is a key component in the analysis (Li et al., 2018; Liu et al., 2021). We here demonstrate the complimentary finding that low frequency medial temporal lobe rhythms might alternatively be best enhanced by more global re-referencing methods, where local methods such as Laplacian reduce power amplitude.

The power results fit one account. On macro contacts, the methods that subtract most of the signal shared between neighbouring contacts, the bipolar and Laplacian references, left the least theta power and the lowest aperiodic background, and the methods that subtract a shaft average or one shared component, AV, MD, GS and ICA, left the most. Theta in the medial temporal lobe is expected to be shared across neighbouring contacts (Buzsáki & Draguhn, 2004), and a reference that removes shared signal removes part of it. For a study of theta on macro contacts a local reference therefore works against the signal of interest, and it is under the local references that theta looks weakest. On microwires the same operations behave differently. The wires cannot be ordered spatially, so a bipolar or Laplacian subtraction removes an arbitrary other wire rather than a neighbour that carries the same theta, and it acts as a monopolar reference. This is why these two methods left power and background on microwires second only to ICA, whereas the average-based methods, which subtract a bundle mean that is nearly the signal itself, left the least. ICA, which removes only the one component with the most uniform weights across a shaft or bundle and returns the rest, subtracted the least shared signal of all methods and left the most theta on both recording types. Whether a method reduces or preserves theta is thus a property of the recording geometry as much as of the method.

Spike–LFP coupling was the least sensitive to the reference. In the hippocampal channels no method differed from another in coupling across all trials, apart from one microwire comparison at 8–12 Hz, and none differed in any coupling memory effect. The same holds for the power memory effects. Two laboratories analysing these recordings under different references would therefore report different theta power, a different background and a different number of retained trials, but neither would find a memory effect that differs between the references, and neither would find one reversed. Which of the small effects passes the threshold under a given reference varies, the retrieval alpha effect was supported under ICA and GS and a trend under AV and MD, but the estimates themselves did not differ between the references. The reference decides how the signal is described, not, whether a memory effect is found.

The aperiodic background deserves separate attention, for two reasons. It carried the most consistent method effects across the two recording types, and it is where the participants differed most: in the all-trial background models the participant term accounted for up to three quarters of the variance, whereas in the memory-effect models of power and coupling it accounted for almost none (Section 3.3). Comparisons of the background between methods therefore depend on treating channels from one participant as related, and a test that treats them as independent would overstate the differences, whereas the memory effects do not depend on this. Because the background moves together with the theta power above it, corrected theta power under one reference cannot be compared with that under another without the background, and we suggest that the fitted background be reported alongside corrected power whenever references differ between studies.

Every memory effect that differed from zero in the hippocampal recordings was negative: power was lower for hits than for misses at retrieval, in the alpha band on macro contacts and in the 4–8 Hz band on microwires, and the one coupling estimate that differed from zero was likewise lower for hits. No positive theta effect was found in power or in spike–LFP coupling. Three features of the analysis bear on this. First, power was read against the fitted aperiodic background, so a memory effect that consists of a broadband shift of the spectrum, which Herweg et al. (2020) identify as one component of reported theta effects, appears in the background parameters rather than in the corrected power. The steeper background for hits than for misses at retrieval reported above, each estimate dependent on one participant. Second, spike–LFP coupling was computed for every spike cluster without selecting neurons by their task response. If only the neurons engaged by the task increase their coupling while others decrease it or shift their preferred phase, the average across all clusters need not rise for hits. Third, in this task successful encoding was accompanied by phase coupling of neurons to faster theta and gamma oscillations, whereas coupling to slower oscillations in the same bands accompanied memory failure (Roux et al., 2022). If successful encoding shifts the frequency at which neurons lock to the LFP rather than strengthening the locking, then a coupling value averaged over a band does not change and the memory effect is invisible to this measure.

Three limits bound these conclusions. First, the comparisons that failed to separate methods are informative only to the extent that the design could have separated them: no spike–LFP pair was complete across all methods and bands, so the coupling null is a statement about the available pairs rather than about every pair that could have been formed. Second, a small number of supported results depend on a single participant, and each is identified where it is reported. Third, the supplementary analysis of all eligible medial temporal channels supports four omnibus tests and one difference between methods in a memory effect that the hippocampal analysis does not (Section 3.3). The boundary of the channel selection, not only the methods themselves, therefore decides which differences reach significance.

In sum, the choice of re-referencing methodology should be an active one that is dependent chiefly on one’s hypothesis and the data at hand.

## Supplementary Materials

### Macro contact spike field coupling

**Figure S1:**
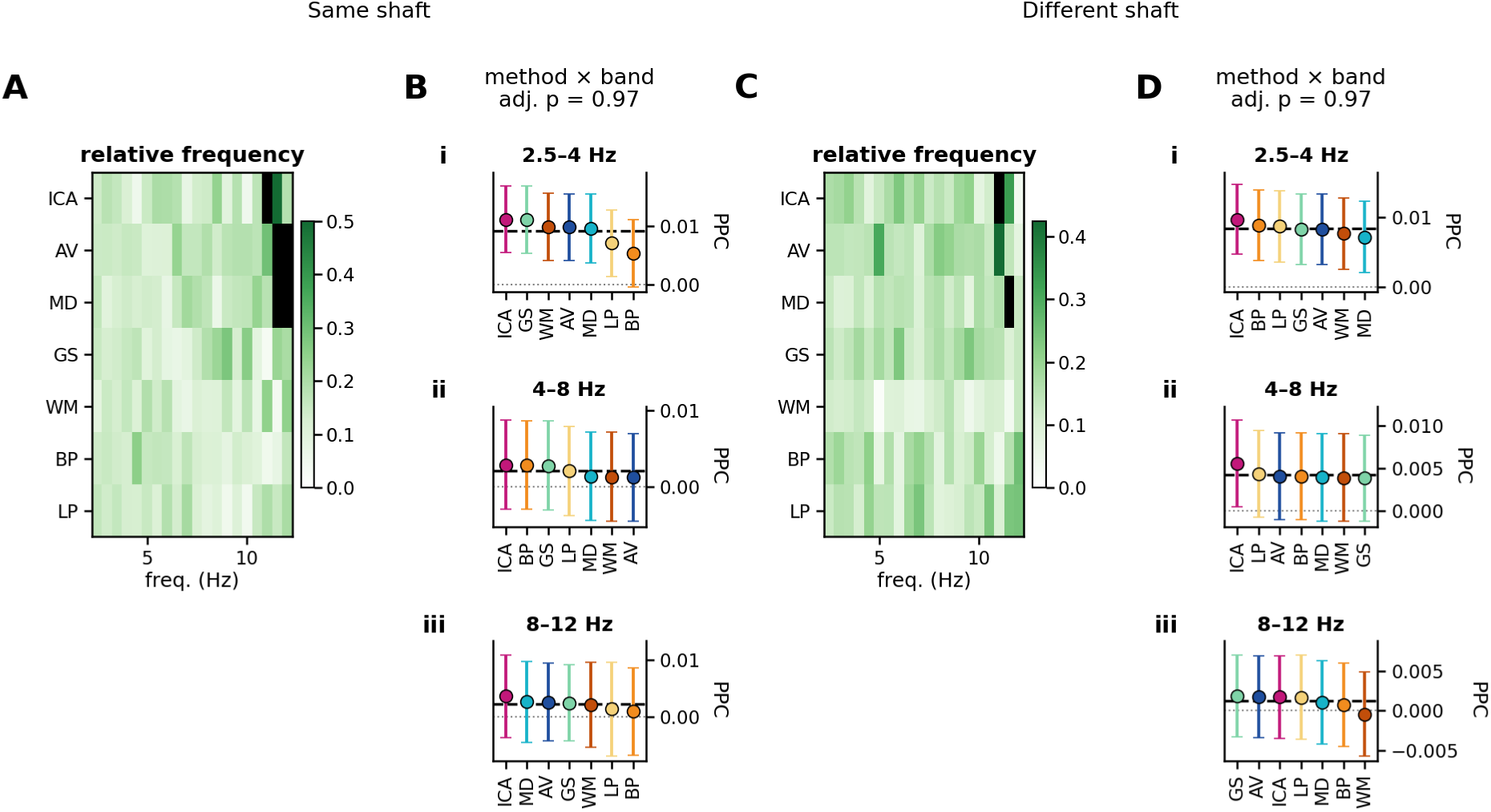
Spike–LFP coupling for all eligible macro-contact LFP channels — every medial temporal channel rather than the ITPC-responsive hippocampal subset used in the main text — across seven re-referencing methods, all trials. A and C show, for same-shaft and different-shaft pairings, each method’s share of the finite coupling estimates at each frequency, black where a method yielded none; these panels are descriptive. B and D show coupling in 2.5–4, 4–8 and 8–12 Hz bands as model-estimated means in PPC units with pointwise 95% confidence intervals, methods rank-ordered by estimate, with brackets for pairwise comparisons passing Benjamini–Hochberg within the panel’s family and the method × band interaction above the band panels. The unit of analysis is the individual spike–LFP pair; counts per panel are in Supplementary Table S1. The correction families are the same as in the main analysis.

### Micro-wire spike field coupling

**Figure S2:**
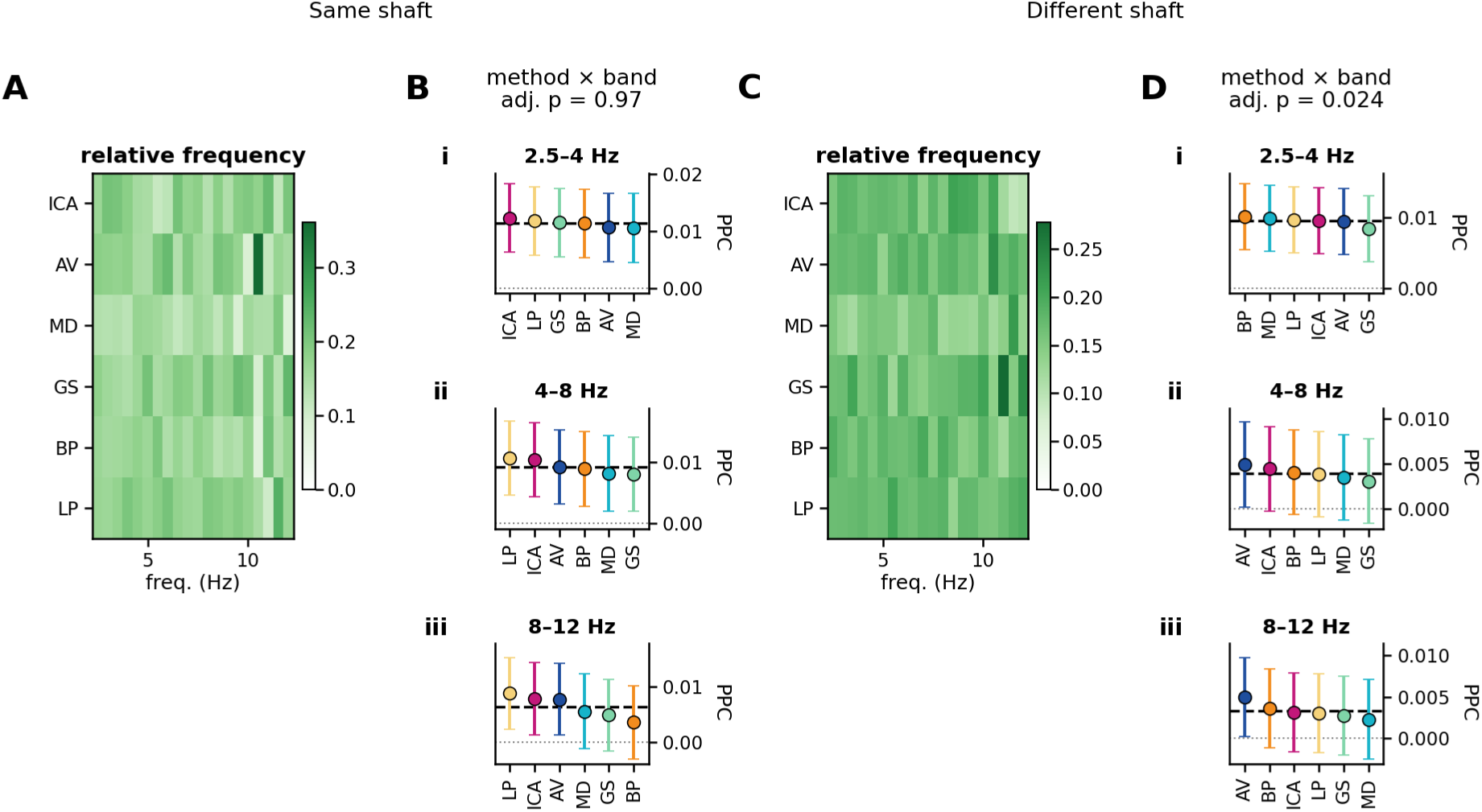
Spike–LFP coupling for all eligible microwire LFP channels — every medial temporal channel rather than the ITPC-responsive hippocampal subset used in the main text — across six re-referencing methods, all trials. A and C show, for same-shaft pairings that exclude the exact same microwire and for different-shaft pairings, each method’s share of the finite coupling estimates at each frequency, black where a method yielded none; these panels are descriptive. B and D show coupling in 2.5–4, 4–8 and 8–12 Hz bands as model-estimated means in PPC units with pointwise 95% confidence intervals, methods rank-ordered by estimate, with brackets for pairwise comparisons passing Benjamini–Hochberg within the panel’s family and the method × band interaction above the band panels. The unit of analysis is the individual spike–LFP pair; counts per panel are in Supplementary Table S1. The correction families are the same as in the main analysis.

### Aperiodic background

**Figure S3:**
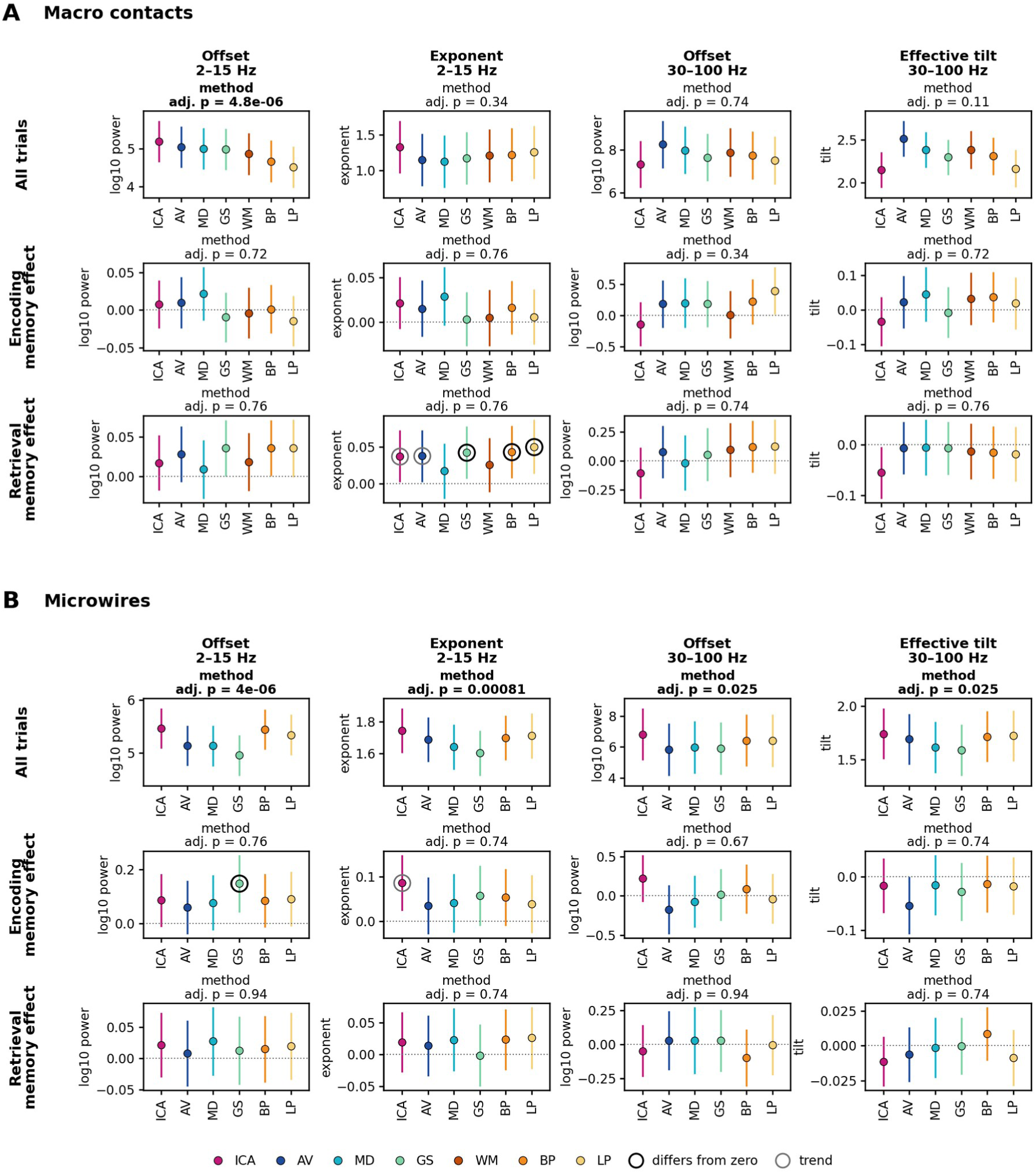
The fitted aperiodic background by re-referencing method for ITPC-responsive hippocampal macro-contact and microwire channels. A, macro contacts; B, microwires. Columns are the four background parameters: the offset and exponent of the 2–15 Hz fit and the offset and effective tilt of the 30 –100 Hz fit (Methods). Rows are all trials, the encoding memory effect and the retrieval memory effect; a memory effect is the difference in the parameter between the hit and miss fits. Points are model-estimated means per method with pointwise 95% confidence intervals; offsets are in log₁₀ power, the exponent and effective tilt are dimensionless. Each panel title gives the method effect after Benjamini–Hochberg across the twenty-four aperiodic tests, in bold where supported. A black ring marks a method whose memory effect differs from zero after Benjamini–Hochberg within the panel’s family, a grey ring a trend. Pairwise comparisons between methods are not drawn; they are in Supplementary Table S4. The unit of analysis is the individual channel; counts per panel are in Supplementary Table S1.

### All eligible medial temporal channels

**Figure S4:**
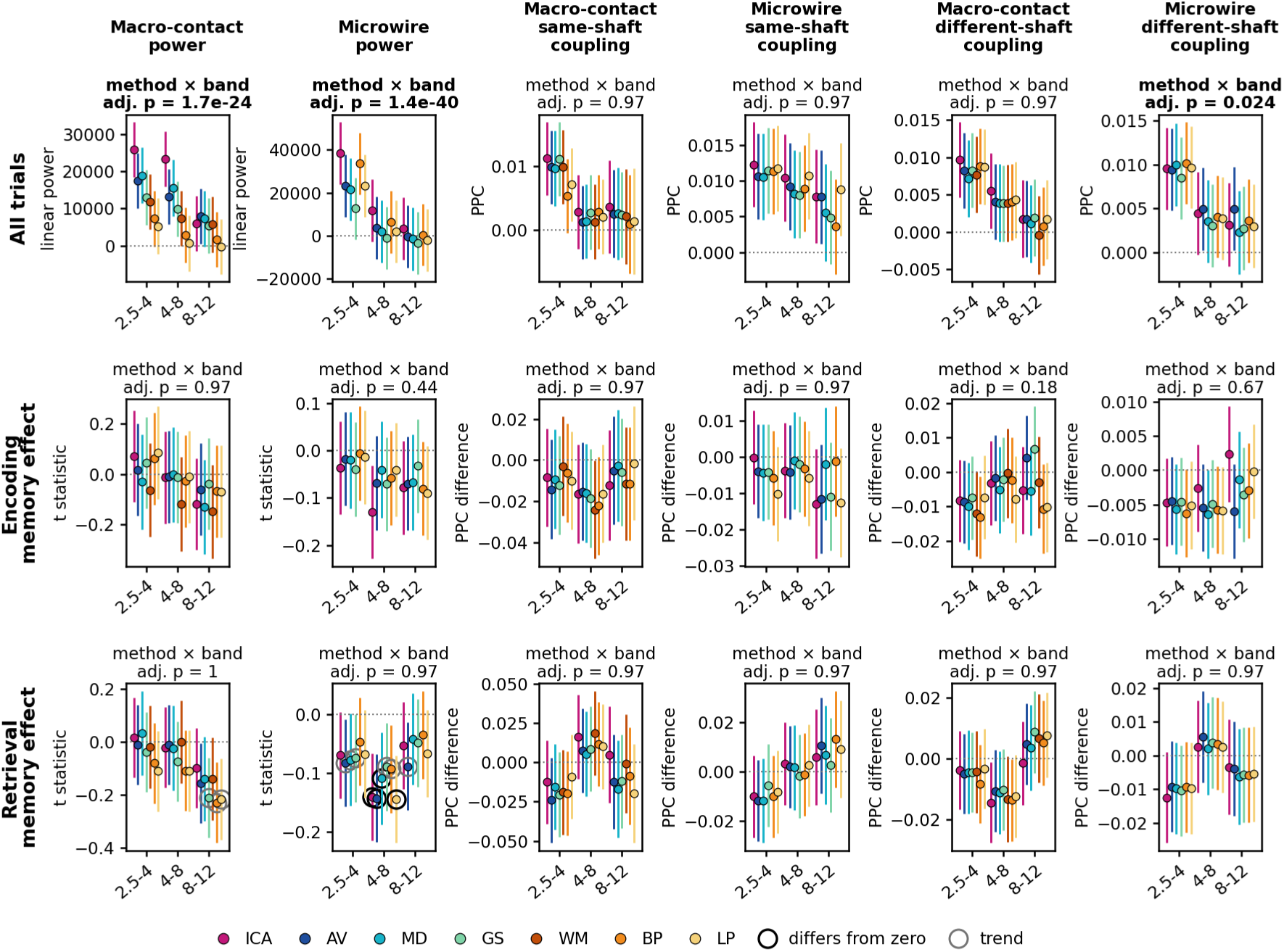
Summary of re-referencing effects across the power and coupling outcomes for all eligible medial temporal channels rather than the ITPC-responsive hippocampal population of the main text. Rows are all trials, the encoding memory effect and the retrieval memory effect; columns are macro-contact power, microwire power, macro-contact and microwire same-shaft coupling, and macro-contact and microwire different-shaft coupling. Within each panel the horizontal axis is the frequency band and colour is the re-referencing method, as elsewhere in the paper; points are model-estimated means with pointwise 95% confidence intervals. Units are linear power for all-trial power, the within-channel hit-versus-miss t statistic for power memory effects, and PPC or the hit-minus-miss PPC difference for coupling. A panel title gives its method × band interaction after Benjamini–Hochberg across the eighteen such tests, in bold where supported. A ring marks a method whose memory effect differs from zero (grey, a trend). Pairwise comparisons between methods are not drawn here; they are in Supplementary Table S4. The unit of analysis is the individual channel for power and the individual spike–LFP pair for coupling, with counts per panel in Supplementary Table S1. No reversal between methods was supported in any panel.

**Figure S5:**
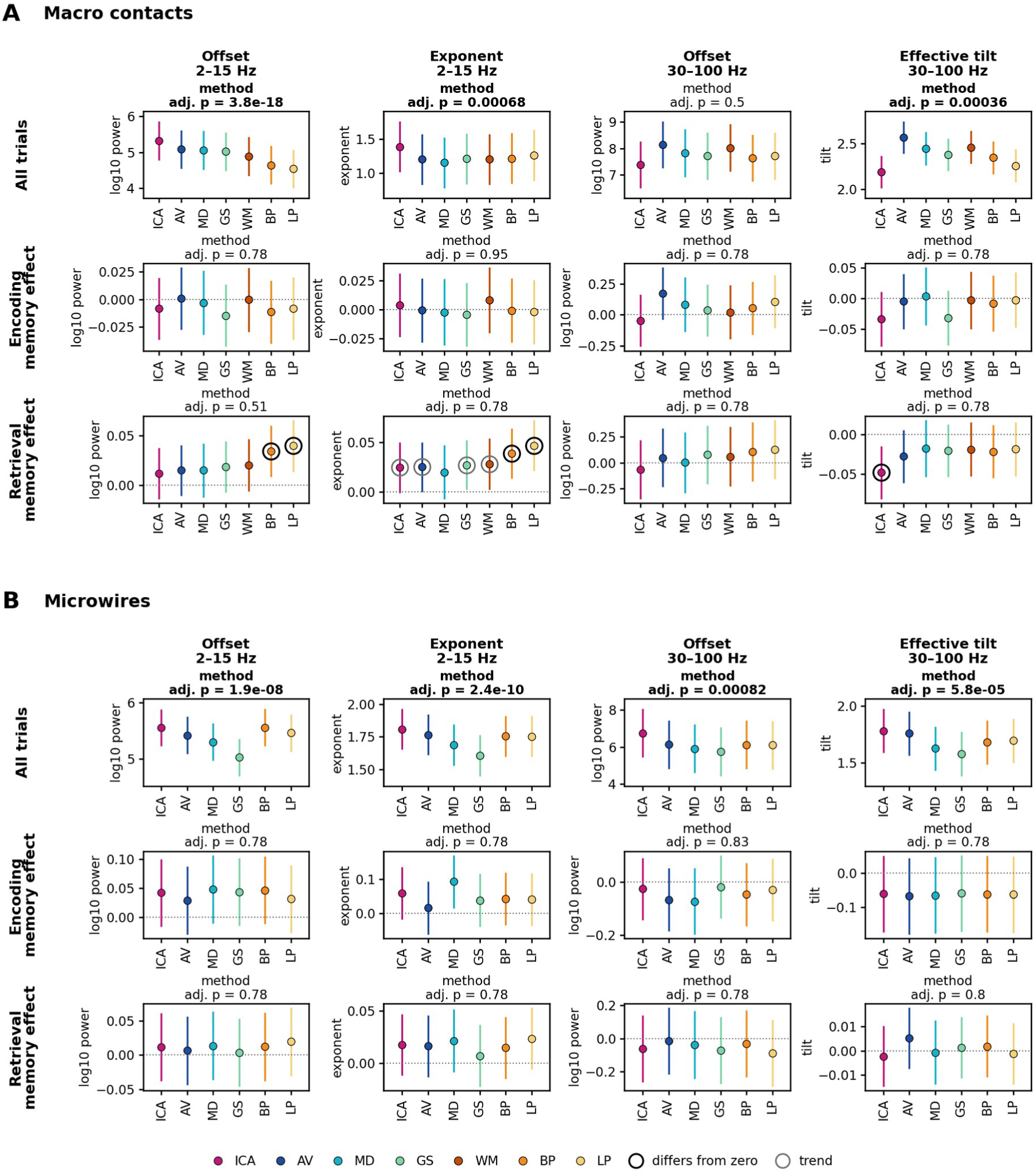
The fitted aperiodic background by re-referencing method for all eligible medial temporal channels rather than the ITPC-responsive hippocampal subset of Supplementary Figure S3. Layout, units, intervals, markers and correction family are as in Supplementary Figure S3; counts per panel are in Supplementary Table S1.

## Supplementary tables

Supplementary Tables S1–S8 are provided as a spreadsheet, one sheet per table, and Table S2 is also reproduced below. Each covers both populations — the ITPC-responsive hippocampal channels of the main text and all eligible medial temporal channels — and both analysis sets: primary, and sensitivity, which additionally excludes aperiodic fits whose error exceeded 0.15 or whose R² fell below 0.85. Adjusted p values are Benjamini–Hochberg-adjusted within the families described in Methods; shading marks adjusted p ≤ 0.05, ≤ 0.01 and < 0.001, and values below 0.001 are shown as <0.001.

Table S1: Numbers of channels (power and aperiodic outcomes) or spike–LFP pairs (coupling), participants and sessions in each model cell, by population, analysis set, recording type, outcome, stage, method and band.

**Table S2:** The primary test of each model — the method × band interaction for the eighteen power and coupling panels and the method effect for the twenty-four aperiodic panels — with F, Satterthwaite degrees of freedom, raw and adjusted p, and the numbers of channels or pairs and participants. The table below shows the adjusted p of each test, shaded as in the spreadsheet, with the numbers of channels or pairs and participants; F, degrees of freedom and raw p are in the spreadsheet.

| Recording | Outcome (test) | All trials |  |  | Encoding memory effect |  |  | Retrieval memory effect |  |  |
| --- | --- | --- | --- | --- | --- | --- | --- | --- | --- | --- |
|  |  | adj. p | n | participants | adj. p | n | participants | adj. p | n | participants |
| ITPC-responsive hippocampal channels (main text) |  |  |  |  |  |  |  |  |  |  |
| **Macro contacts** | Corrected power (method × band) | 0.288 | 166 | 15 | 0.931 | 130 | 14 | 0.931 | 129 | 15 |
| | Same-shaft coupling (method × band) | 0.931 | 506 | 13 | 0.931 | 369 | 13 | 0.767 | 368 | 13 |
| | Different-shaft coupling (method × band) | 0.931 | 1,664 | 11 | 0.931 | 1,377 | 11 | 0.906 | 1,384 | 11 |
|  | Offset 2–15 Hz (method) | <0.001 | 177 | 15 | 0.725 | 130 | 14 | 0.762 | 129 | 15 |
|  | Exponent 2–15 Hz (method) | 0.337 | 177 | 15 | 0.762 | 130 | 14 | 0.762 | 129 | 15 |
|  | Offset 30–100 Hz (method) | 0.740 | 177 | 15 | 0.337 | 130 | 14 | 0.741 | 129 | 15 |
|  | Effective tilt 30–100 Hz (method) | 0.114 | 177 | 15 | 0.725 | 130 | 14 | 0.762 | 129 | 15 |
| **Microwires** | Corrected power (method × band) | <0.001 | 429 | 16 | 0.767 | 318 | 14 | 0.931 | 299 | 16 |
| | Same-shaft coupling (method × band) | 0.906 | 982 | 15 | 0.767 | 770 | 14 | 0.671 | 760 | 14 |
| | Different-shaft coupling (method × band) | 0.967 | 3,998 | 12 | 0.931 | 3,032 | 10 | 0.931 | 2,978 | 12 |
|  | Offset 2–15 Hz (method) | <0.001 | 446 | 16 | 0.762 | 318 | 14 | 0.937 | 299 | 16 |
|  | Exponent 2–15 Hz (method) | <0.001 | 446 | 16 | 0.740 | 318 | 14 | 0.740 | 299 | 16 |
|  | Offset 30–100 Hz (method) | 0.025 | 446 | 16 | 0.667 | 318 | 14 | 0.937 | 299 | 16 |
|  | Effective tilt 30–100 Hz (method) | 0.025 | 446 | 16 | 0.740 | 318 | 14 | 0.740 | 299 | 16 |
|  |  | adj. p | n | participants | adj. p | n | participants | adj. p | n | participants |
| All eligible medial temporal channels (supplementary population) |  |  |  |  |  |  |  |  |  |  |
| Macro contacts | Corrected power (method × band) | <0.001 | 404 | 15 | 0.969 | 333 | 15 | 0.998 | 332 | 15 |
|  | Same-shaft coupling (method × band) | 0.969 | 1,003 | 13 | 0.969 | 806 | 13 | 0.969 | 802 | 13 |
|  | Different-shaft coupling (method × band) | 0.969 | 4,324 | 14 | 0.176 | 3,782 | 14 | 0.969 | 3,786 | 14 |
|  | Offset 2–15 Hz (method) | <0.001 | 417 | 15 | 0.775 | 333 | 15 | 0.515 | 332 | 15 |
|  | Exponent 2–15 Hz (method) | <0.001 | 417 | 15 | 0.955 | 333 | 15 | 0.775 | 332 | 15 |
|  | Offset 30–100 Hz (method) | 0.496 | 417 | 15 | 0.775 | 333 | 15 | 0.775 | 332 | 15 |
|  | Effective tilt 30–100 Hz (method) | <0.001 | 417 | 15 | 0.775 | 333 | 15 | 0.777 | 332 | 15 |
| Microwires | Corrected power (method × band) | <0.001 | 1,553 | 16 | 0.435 | 1,266 | 16 | 0.969 | 1,232 | 16 |
|  | Same-shaft coupling (method × band) | 0.969 | 3,453 | 16 | 0.969 | 2,816 | 16 | 0.969 | 2,738 | 16 |
|  | Different-shaft coupling (method × band) | 0.024 | 16,372 | 15 | 0.669 | 13,938 | 15 | 0.969 | 13,685 | 15 |
|  | Offset 2–15 Hz (method) | <0.001 | 1,608 | 16 | 0.775 | 1,266 | 16 | 0.775 | 1,232 | 16 |
|  | Exponent 2–15 Hz (method) | <0.001 | 1,608 | 16 | 0.775 | 1,266 | 16 | 0.775 | 1,232 | 16 |
|  | Offset 30–100 Hz (method) | <0.001 | 1,608 | 16 | 0.833 | 1,266 | 16 | 0.775 | 1,232 | 16 |
|  | Effective tilt 30–100 Hz (method) | <0.001 | 1,608 | 16 | 0.775 | 1,266 | 16 | 0.800 | 1,232 | 16 |

Table S3: Model-estimated mean of every method in every band, in the outcome’s units, with pointwise 95% confidence intervals. Memory-effect means are also tested against zero, with adjustment across the tests of the same panel; all-trial means are descriptive.

Table S4: Every planned within-band pairwise comparison between methods: the difference (method A minus method B) with its pointwise 95% confidence interval, F, Satterthwaite degrees of freedom, raw p and p adjusted across the comparisons of the same panel, and the channels or pairs and participants the two methods share.

Table S5: Reversal classification. Every planned pairwise comparison of memory effects with its estimate and adjusted p, the two methods’ memory effects with their adjusted p against zero, and the resulting classification. A reversal was counted only when the two memory effects had opposite signs and both, together with their direct comparison, met the adjusted criterion; no comparison met it, in either population or analysis set.

Table S6: Random intercepts in each final model, and the optional intercepts removed under the prespecified rule because their variance was estimated at zero, with the numbers of observations, channels or pairs, and participants. Participant and channel or pair intercepts were never removed.

Table S7: Every effect that survived adjustment — omnibus tests, pairwise comparisons and tests against zero — re-estimated with each participant removed in turn: the estimate with all participants (F for omnibus tests) and its adjusted p, and, for each removal, the re-estimate and its unadjusted p. A result depends on a single participant when that p exceeds 0.05 under at least one removal. Re-estimates are given to four significant digits.

Table S8: Variance of each random intercept, and of the residual, as a proportion of the total variance of each model; the proportions in a row sum to one. “Removed” marks an optional intercept dropped because its variance was estimated at zero (Table S6); a dash marks an intercept that is not part of that model.

